# Combined protein and transcript single cell RNA sequencing in human peripheral blood mononuclear cells

**DOI:** 10.1101/2020.09.10.292086

**Authors:** Jenifer Vallejo, Ryosuke Saigusa, Rishab Gulati, Yanal Ghosheh, Christopher P. Durant, Payel Roy, Erik Ehinger, Vasantika Suryawanshi, Tanyaporn Pattarabanjird, Lindsey E. Padgett, Claire E. Olingy, David B. Hanna, Alan L. Landay, Russell P. Tracy, Jason M. Lazar, Wendy J. Mack, Kathleen M. Weber, Adaora A. Adimora, Howard N. Hodis, Phyllis C. Tien, Igho Ofotokun, Sonya L. Heath, Rafael Blanco-Dominguez, Huy Q. Dinh, Avishai Shemesh, Coleen A. McNamara, Lewis L. Lanier, Catherine C. Hedrick, Robert C. Kaplan, Klaus Ley

## Abstract

Cryopreserved peripheral blood mononuclear cells (PBMCs) are frequently collected and provide disease- and treatment-relevant data in clinical studies. Here, we developed combined protein (40 antibodies) and transcript single cell (sc)RNA sequencing in PBMCs. Among 31 participants in the WIHS Study, we sequenced 41,611 cells. Using Boolean gating followed by Seurat UMAPs and Louvain clustering, we identified 58 subsets among CD4 T, CD8 T, B, NK cells and monocytes. This resolution was superior to flow cytometry, mass cytometry or scRNA-sequencing without antibodies. Since the transcriptome was not needed for cell identification, combined protein and transcript scRNA-Seq allowed for the assessment of disease-related changes in transcriptomes and cell type proportion. As a proof-of-concept, we showed such differences between healthy and matched individuals living with HIV with and without cardiovascular disease. In conclusion, combined protein and transcript scRNA sequencing is a suitable and powerful method for clinical investigations using PBMCs.

## Introduction

PBMCs are a rich source of disease- and treatment-relevant information.[3, 9, 22, 30, 38, 53, 60–62] PBMCs can be analyzed without mechanical or enzymatic dissociation, which are known to alter cell surface markers and transcriptomes.[56] PBMCs can be cryopreserved without loss of viability. At the most basic level, lymphocytes and monocyte can be distinguished by morphology using automated cell counters (CBC).[4] Current practice is to use flow cytometry of between 8 and 16 markers.[40, 43, 52] More recently, mass cytometry became available,[1, 11, 47, 57] allowing for analysis of up to 40 markers. Single cell RNA-sequencing (scRNA-Seq) allows the interrogation of expressed genes[8, 28, 35, 48, 63] and surface markers.[48, 64]

In immune cells, the correlation between mRNA and surface expression of any given surface marker is weak.[25] This is because cell surface expression is not only determined by gene expression, but also by posttranslational protein modifications,[26] trafficking to the cell surface, protein stability, and proteolytic modifications. Cell types in PBMCs have been defined by flow cytometry, and the surface markers of the major cell types are very well known. Yet, it is difficult to call even major cell types by scRNA-Seq. For example, CD4 T cells are not resolved from CD8 T cells and natural killer (NK) cells.[59] To capitalize on the vast flow and mass cytometry literature, it is necessary to assess cell surface phenotype along with transcriptomes.

Only two publications reported single cell transcriptomes from patients with atherosclerosis (carotid endarterectomy specimens and matched PBMCs).[9, 58] 1,652 PBMCs from one patient were analyzed by 10x Genomics 3’ and cellular indexing of transcriptional epitope sequencing (CITE-Seq),[35, 48] using a panel of 21 antibodies. No healthy control PBMCs were studied. A recent study reported the effect of HIV infection on PBMC transcriptomes,[20] focusing on acute HIV infection (before antiretroviral therapy started) and reporting PBMC transcriptomes in four patients at 8 defined time points (average of 1,976 PBMC transcriptomes per participant and condition). No scRNA-Seq or CITE-Seq studies of PBMCs of people living with chronic HIV infection have been reported. No single cell studies of the interaction between HIV and CVD are available.

Here, we report transcriptomes and cell surface phenotypes of almost 42,000 PBMCs using the targeted scRNA-Seq BD Rhapsody platform[8, 28] that simultaneously provides single cell surface phenotype (40 monoclonal antibodies, mAbs) and transcriptomes (485 immune and inflammatory transcripts) in the same cells. As a proof-of-concept, we show significant differences in cell proportions and cell transcriptomes between healthy subjects and matched subjects living with HIV or cardiovascular disease from the WIHS cardiovascular sub-study. WIHS is an ongoing multi-center, prospective, observational cohort study of women with or at risk of HIV infection. PBMCs were cryopreserved on liquid N2, following strict standard operating procedures that ensured preservation of cell surface phenotype, viability, and transcriptomes.

## Material and Methods

### Study characteristics and sample selection

The Women’s Interagency HIV Study (WIHS) was initiated in 1994 at six (now expanded to ten) U.S. locations.[13, 16] It is an ongoing prospective study of over 4,000 women with or at risk of HIV infection. Recruitment in the WIHS occurred in four phases (1994-1995, 2001-2002, 2010-2012, and 2013-2015) from HIV primary care clinics, hospital-based programs, and community outreach and support groups. Briefly, the WIHS involves semi-annual follow-up visits, during which participants undergo similar detailed examinations, specimen collection, and structured interviews assessing health behaviors, medical history, and medication use. All participants provided informed consent, and each site’s Institutional Review Board approved the studies.

All participants in the current analysis were part of a vascular sub-study nested within the WIHS.[13, 16, 18] The baseline visit for the vascular sub-study occurred between 2004 and 2006, and a follow-up visit occurred on average seven years later. Participants underwent high-resolution B-mode carotid artery ultrasound to image six locations in the right carotid artery: the near and far walls of the common carotid artery, carotid bifurcation, and internal carotid artery. A standardized protocol was used at all sites,[18] and measurements of carotid artery focal plaque, a marker of subclinical atherosclerosis, were obtained at a centralized reading center (U. of Southern California). Subclinical CVD (sCVD) was defined based on the presence of one or more carotid artery lesions.[18]

From the initial 1,865 participants in the WIHS vascular sub-study, 32 participants were selected for scRNA-seq analysis. CVD was defined as presence of carotid artery focal plaque at either vascular sub-study visit to define four groups of eight participants each: HIV-, HIV+CVD-, HIV+CVD+, HIV+CVD+ on CRT. Because we were interested in the joint relationships of HIV infection and sCVD with surface marker and RNA expression by different cell subtypes, we selected matched samples based on HIV, CVD and cholesterol-reducing treatment (CRT, mostly statins). The latter was done because we found that CRT had a major impact on monocyte transcriptomes.[7]. HIV infection status was ascertained by enzyme-linked immunosorbent assay (ELISA) and confirmed by Western blot. Non-CVD participants with self-reported coronary heart disease or current lipid-lowering therapy use were excluded. Participants were formed in quartets matched by race/ethnicity (except one quartet), age (± 5 years) at the baseline vascular sub-study (except one quartet where the age difference was more but all the women were post-menopausal), visit number, smoking history, and date of specimen collection (within 1 year).

Demographic, clinical, and laboratory variables were assessed from the same study visit using standardized protocols. The median age at the baseline study visit was 55 years, and 96% of participants were either of Black race or Hispanic ethnicity. Most (86%) reported a history of smoking. Substance use was highly prevalent, with 43% of HIV+ and 50% of HIV- participants reporting either a history of injection drug use; current use of crack, cocaine, or heroin; or alcohol use (≥14 drinks per week). Among HIV+ participants, over 80% reported use of HAART at the time PBMCs were obtained, and 59% reported an undetectable HIV-1 RNA level. The median CD4+ T-cell count was 585 cells/μL (IQR 382-816) in HIV+ women without sCVD and 535 cells/μL (IQR 265-792) in HIV+ women with sCVD.

### Preparation of PBMC samples for combined protein and RNA-seq

To avoid batch effects, sixteen samples each were processed on the same day. PBMC tubes were thawed in a 37°C water bath and tubes filled with 8 mL of complete RPMI-1640 solution (**Table S1**; cRPMI-1640 contains human serum albumin, HEPES, sodium pyruvate, MEM-NEAA, penicillin-streptomicyn, GlutaMax, and mercaptoethanol). The tubes were centrifuged at 400 xg for 5 minutes and pellets resuspended in cold staining buffer (2 % fetal bovine serum (FBS) in phosphate-buffered saline (PBS)). All reagents, manufacturers, and catalogue numbers are listed in **Table S1**. Manual cell counting was performed by diluting cell concentration to achieve 100-400 cells per hemocytometer count. Cells were aliquoted to a count of 1 million cells each and incubated on ice with Fc Block (BD, **Table S1**) at a 1:20 dilution, centrifuged at 400 xg for 5 minutes, resuspended in 180 μL of SB and transferred to their respective sample multiplexing kit tubes (BD). The cells were incubated for 20 minutes at room temperature, transferred to 5 mL polystyrene tubes, washed 3 times and centrifuged at 400 xg for 5 minutes. The cells were resuspended in 400 μL of staining buffer and 2 μL of DRAQ7 and calcein AM were added to each tube. The viability and cell count of each tube was determined using the BD Rhapsody scanner (**Table S2**). Tube contents were pooled in equal proportions with total cell counts not to exceed 1 million cells. The tubes were then centrifuged at 400 xg for 5 minutes and resuspended in a cocktail of 40 AbSeq (**Table S3**) antibodies (2 μL each and 20 μL of staining buffer) on ice for 30-60 minutes per manufacturer’s recommendations. The tubes were then washed with 2 mL of SB followed by centrifugation at 400 xg for 5 minutes. This was repeated two more times for a total of 3 washes. The cells were then counted again using the scanner.

### Library preparation

Cells were loaded at 800-1000 cells/μL into the primed plate per the BD user guide. The beads were isolated with a magnet and the supernatant removed. Reverse transcription was performed at 37 °C on a thermomixer at 1200 rpm for 20 minutes. Exonuclease I was incubated at 37 °C on a thermomixer at 1200 rpm for 30 minutes and then immediately placed on a heat block at 80 °C for 20 minutes. The tube was placed on ice followed by supernatant removal while beads were on a magnet. The beads were resuspended in BD bead resuspension solution. Then, the tubes were stored at 4 °C until further processing. Per BD’s protocol, the reagents for PCR1 including the BD Human Immune Response Panel and a custom panel of ~100 genes (**Table S4**) were added to the beads. Samples were aliquoted into four 0.2 mL strip PCR tubes and incubated for 10 cycles according to BD’s protocol for PCR1. A double size selection was performed to remove high genomic DNA fragments by adding 0.7x volume AMPure XP SPRI beads to the PCR products. After incubation, the supernatant is recovered and transferred to a new tube followed by purifying the supernatant with an additional 100 μL of AMPure XP beads (sample tags and antibodies). The RNA tube was washed twice with 500 μL of 80 % ethanol. 550 μl of supernatant were removed from the antibody tube followed by two washes with 500 μL of 80 % ethanol. The cDNA was eluted off the beads using 30 μL of BD elution buffer and then transferred to a 1.5 mL tube.

### Pre-sequencing quality control (QC)

QC/ and quantification was performed on the tube containing AbSeq and Sample Tags using Agilent TapeStation high sensitivity D1000 screentape. 5 μL from each tube (mRNA and Ab/ST) was then added to their respective tubes containing the reagents for PCR2. Each tube had 12 cycles of PCR performed according to BD’s user guide. Each tube was cleaned with AMPure XP beads at 0.8X for mRNA and 1.2X for sample tags. Two 200 μL washes were performed during the clean-up using 80 % ethanol per sample. The cDNA was eluted off using BD elution buffer. QC/ and quantification was performed using Agilent TapeStation high sensitivity D1000 screen tape and Qubit double stranded high sensitivity DNA test kit. The mRNA was then diluted, if necessary, to a concentration of 1.2-2.7 ng/μL and the antibody and sample tag libraries from PCR2 were diluted, if needed, to a concentration of 0.5-1.1 ng/μL. From each sample 3 μL were added to a volume of 47 μL of reagents for PCR3 as described by BD’s user guide following the protocol and number of cycles listed, except for AbSeq, which had 9 cycles of PCR performed as determined by previous optimization. The three libraries were then cleaned with AMPure XP beads at 0.7X for AbSeq and 0.8X for sample tags. Samples were washed twice with 200 μL of 80 % ethanol. The cDNA was eluted off the beads using BD’s elution buffer. Final QC and quantification was performed using TapeStation and Qubit kits and reagents.

### Sequencing

The samples were pooled and sequenced to the following nominal depth recommended by BD: AbSeq: n x 1000 reads per cell, where n is the plexity of AbSeq used; mRNA: 20,000 reads per cell; Sample Tags: 600 reads per cell. Thus, a total of 60,600 reads per cell were desired for sequencing on the NovaSeq. The samples and specifications for pooling and sequencing depth, along with number of cells loaded onto each plate was optimized for S1 and S2 100 cycle kits (Illumina) with the configuration of 67×8×50 bp. Once sequencing was complete, a FASTA file was generated by BD as a reference for our AbSeq and genes we targeted with these assays. The FASTA file and FASTQ files generated by the NovaSeq were uploaded to Seven Bridges Genomics pipeline, where the data was filtered and matrices and csv files were generated. This analysis generated draft transcriptomes and surface phenotypes of 54,078 cells (496 genes, 40 antibodies). 11 genes were not expressed, leaving 485 genes for analysis.

### Doublet Removal

Based on the 4 sample tags used per plate, 8,359 doublets were removed. The remaining 45,719 cells were analyzed using the Doublet Finder package on R (https://github.com/chris-mcginnis-ucsf/DoubletFinder) with the default doublet formation rate (7.5%). This removed another 3,322 doublets, leaving 42,397 Cells. Finally, we removed all cells that had less than 128 (2^7^) antibody molecules sequenced. This removed 786 noisy cells, resulting in 41,611 cell transcriptomes. All antibody data were CLR (centered log-ratio) normalized and converted to log_2_ scale. All transcripts were normalized by total UMIs in each cell and scaled up to 1000.

### Thresholding

Preliminary experiments showed that each antibody had both specific and non-specific binding, as expected. To remove the non-specific signal, each antibody threshold (**Table S5**) was obtained by determining its expression in a known negative cell. To identify the thresholds, biaxial plots of mutually exclusive markers were used to best separate the positive populations from the noise. In combined protein and transcript panel single cell sequencing, non-specific background staining is caused by incomplete Fc block and oligonucleotide-tagged antibody being trapped in the nanowell.[48] Ridgeline plots of the unthresholded and thresholded antibody expressions for each main cell type are shown in **Supplemental Figure S1**, which indicates how the thresholding worked on each antibody expression.

### Clustering

Clustering was performed using UMAP (Uniform Manifold Approximation and Projection) and Louvain clustering.[50] UMAP is a manifold learning technique for dimensionality reduction. It is based on neighborhood graphs, which captures the local relationship in the data. UMAP is able to maintain local structure and also preserve global distances in the reduced dimension, i.e. the cells that are similar in the high dimension remain close-by in the 2 dimensions and the cells that different are apart in the 2 dimensions. The clustering parameters used were: n_neighbors = 100, n_pcs = 50, min_dist = 1, spread = 1, random state = 42. Louvain resolution was set at 0.8. Subclustering of each major cell type was based on all non-negative antibodies (**Table S6**). Gates were overlaid and used in all subsequent UMAP figures (cell numbers in **Table S7**)

### Cluster Assignment

In CD4 T cells, 4 of the initial clusters were further divided based on the expression of CD11c, CD56, CD25, CD127, CXCR3, and CCR2. CD8 T cells had two clusters that were divided based on CD11c, CD16, and CXCR3 surface marker expression. One cluster from classical monocytes and one cluster from intermediate monocytes were further divided based on CCR7 and CD152 expression, respectively. In non-classical monocytes, one cluster showed differential expression of CD36 and CD152 expression and was divided in two. In B cells, one cluster was split because it showed differential expression of CD25 and CXCR3 within the cluster. Finally, two clusters from NK cells were split due to CD16, CD56, and CD11c expression.

### Comparing Gene Expression among Participant Types

To determine differential expression (DE) among the four types of participants, we use the Seurat package [49] in R with no thresholds over avg_logFC, minimum fraction of cells required in the two populations being compared, minimum number of cells and minimum number of cells expressing a feature in either group. We filtered for adjusted p<0.05 and compared HIV-, HIV+CVD-, HIV+CVD+, and HIV+CVD+CRT+. From this data, volcano plots were generated using ggplot2 and ggrepel packages in R. Axes were restricted to the range of (−2,2) on the x-axis and (0,20) on the y-axis. Genes outside these ranges were bounded to the corresponding limit of the axes. Exact p-values of differential gene expression in all major cell types and the top 10 highly regulated genes for the main cell types are shown in **Table S8** and **S9**, respectively.

### Comparing Cell Proportions

To find changes in proportions, we identified the cell numbers for each participant in each cluster (**Table S10)**. Statistical differences in cell proportions were calculated by log-odds ratio defined as p/(1-p) where p is the proportion of cells, followed by ANOVA and Tukey’s multiple comparison test between the four groups. For clarity, the data are presented as percentage of cells.

### Correlation Analysis

We correlated each antibody to its corresponding gene(s) using Spearman rank correlation and significance (R package). For each combination of gene-antibody, we discarded cells that had values below the corresponding threshold for that antibody as well as cells with zero counts for that gene. After this filter, any gene-antibody combination that had 10 cells or less was deemed insignificant. Finally, all non-significant (p-value > 0.05) were designated a nominal value of zero as the Spearman rank correlation coefficient and we selected only those genes or antibodies that had at least one correlation whose coefficient >= 0.25 or whose coefficient <= −0.25. All significant non-negative correlations are reported in **Table S11**.

## Results

### Identification of main cell types based on antibody expression

To identify the major known cell types, we used biaxial gating on CD3, CD19, CD4, CD8, CD14, CD16, and CD56. This approach defines (**Figure 1A-E**):

- B cells: CD19+ CD3−
- T cells: CD19− CD3+
- CD4 T cells: CD4+ CD8− T cells
- CD8 T cells: CD8+ CD4− T cells
- Monocytes (M): CD19-CD3-CD56−
- Classical (CM): CD14+CD16−
- Intermediate (INT): CD14+CD16+
- Nonclassical (NCM): CD14-CD16+CD56−
- NK cells (NK): CD4− CD56+ CD14− CD20− CD123− CD206−

**Figure 1.**
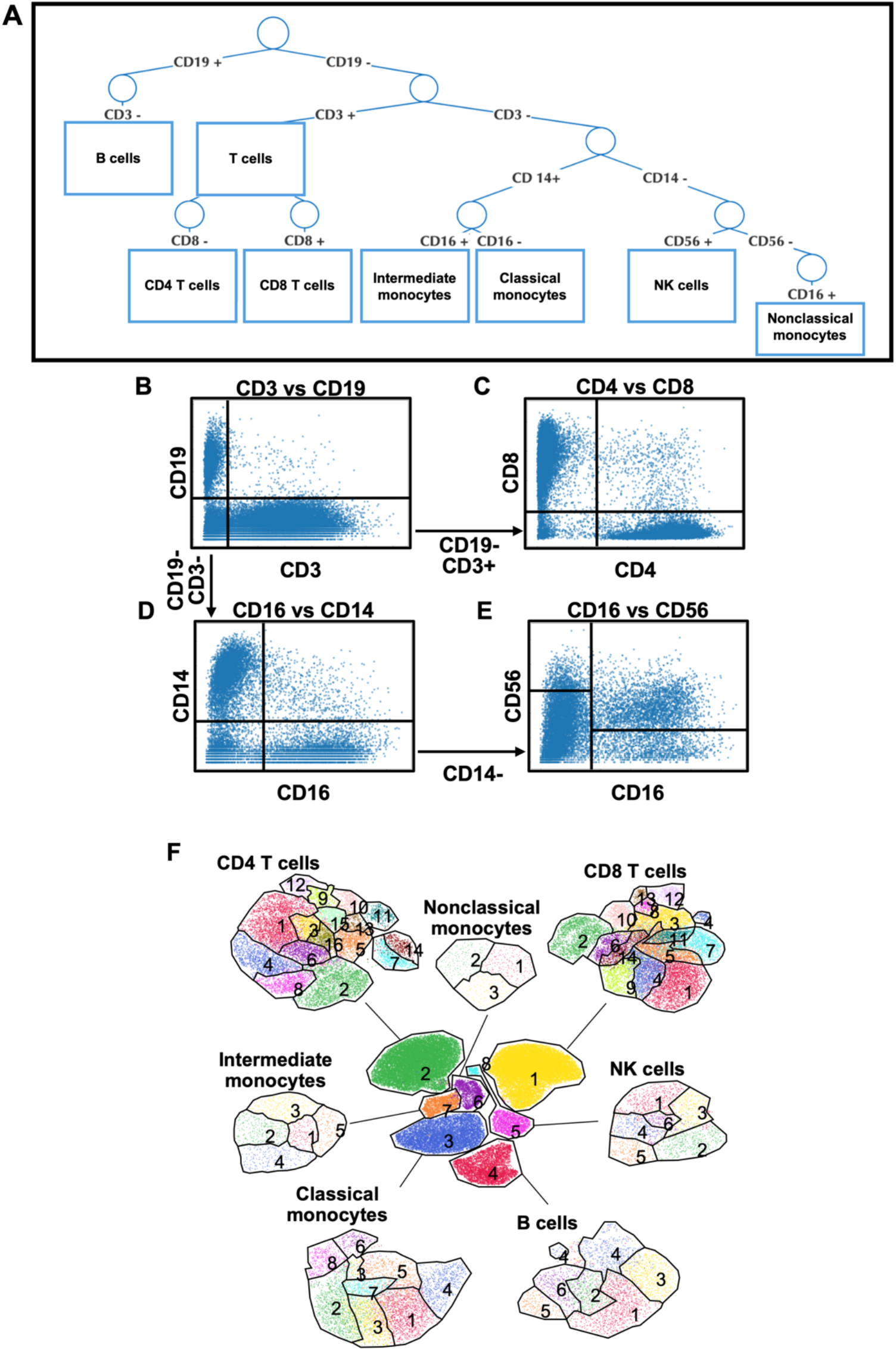
Gating scheme (A), biaxial dot plots (B-E) to identify major known cell types, and (F) antibody-based UMAP clustering of major cell types. PBMCs from 32 WIHS participants were hash-tagged and stained with 40 oligonucleotide-tagged mAbs (**Table S3**). (**B**) B cells were defined as CD19+CD3- and T cells as CD19-CD3+. (**C**) T cells were identified as CD4 (CD4+CD8-) or CD8 (CD4− CD8+). (**D**) All CD19-CD3− cells were gated for CD14 and CD16, with CD14+CD16− cells being classical (CM) and CD14+CD16+ being intermediate (INT) monocytes. (**E**) The CD14-CD16+ cells from panel D contain NK cells, which were identified by CD56 and defined as CD56+CD14-CD20-CD123-CD206-. Most of the remaining CD56− CD16+ cells were nonclassical monocytes (NCM). (**F**) The major known cell types were UMAP-Louvain-clustered by CD3, CD19, CD14, CD16, and CD56 surface expression (central panel). Then, each major known cell type was UMAP-Louvain-clustered by all non-negative surface markers. CD4 T cells formed 16 clusters, cluster numbers indicated; CD8 T cells formed 14 clusters; Classical monocytes (CM) formed 8, Intermediate monocytes (INT) 5, and Nonclassical monocytes (NCM) 3 clusters. B cells and NK cells formed 6 clusters each.

CD3 and CD19 expression are mutually exclusive and specific for T and B cells, respectively. As is standard in the NK cell field,[31] the CD16- immature NK cells were gated based on higher levels of CD56 as shown in **Figure 1E**. The mature NK cells were CD19-CD3-CD16+CD56+. One CD16+CD56- cluster was also identified as NK cells. This resulted in 2,919 B cells, 11,045 CD4 T cells, 12,843 CD8 T cells, 5,145 CM, 1009 INT, 475 NCM and 1,843 NK cells. Each of these major cell types was then re-clustered separately, using Seurat [49] to construct UMAPs with Louvain clustering (**Figure 1F**). Like in flow or mass cytometry, we clustered on antibody staining only. This “preserves” the transcriptomes for investigations into disease- and treatment-related changes. Using this approach, we identified 16 CD4 T cell subsets, 14 CD8 T cell subsets, 8 CM subsets, 3 NCM subsets, 5 INT subsets, 6 B cell subsets and 6 NK cell subsets (**Figure 1F**). The corresponding feature maps are shown in **Figure S2**. Trying to find these cell types based on transcriptomes was unsatisfactory (**Figure S3**).

### Cell subsets calling using 40 surface markers

Next, we constructed heat maps for all antibodies that were significantly differentially expressed in at least one subset (**Figure 2**). This information allowed us to call all CD4 and CD8 T cell subsets in accordance with published immunology work. Among CD4 T cells, CD2 was expressed in almost all cells, as expected. The high affinity IL2 receptor IL2RA (CD25) was expressed in about a third of the CD4 T cells and was strikingly high in cluster 13, which was also low for IL7 receptor (CD127), defining cluster 13 as regulatory T cells (Tregs). CD45RA and RO were mutually exclusive, separating naive and antigen-experienced CD4 T cells. CXCR3 (CD183) identifies T-helper-1 (Th1) cells and was highly expressed in clusters 5, 14, and 16. Cluster 14 co-expressed CXCR5 (CD185) with CXCR3. Cluster 7 expressed CXCR5 as the only chemokine receptor, suggesting it may contain follicular helper (TFH) T cells. Based on surface marker information, all CD4 T cell clusters were called (**Figure 2A**). All CD8 T cells expressed CD2. Cluster 3 exclusively expressed CD9 and CD36, identifying these cells as NK-like CD8 cells. Clusters 7 and 13 were identified as NK-like T cells with a CD45RA+ terminally differentiated memory (EMRA) phenotype (**Figure 2B**).

**Figure 2.**
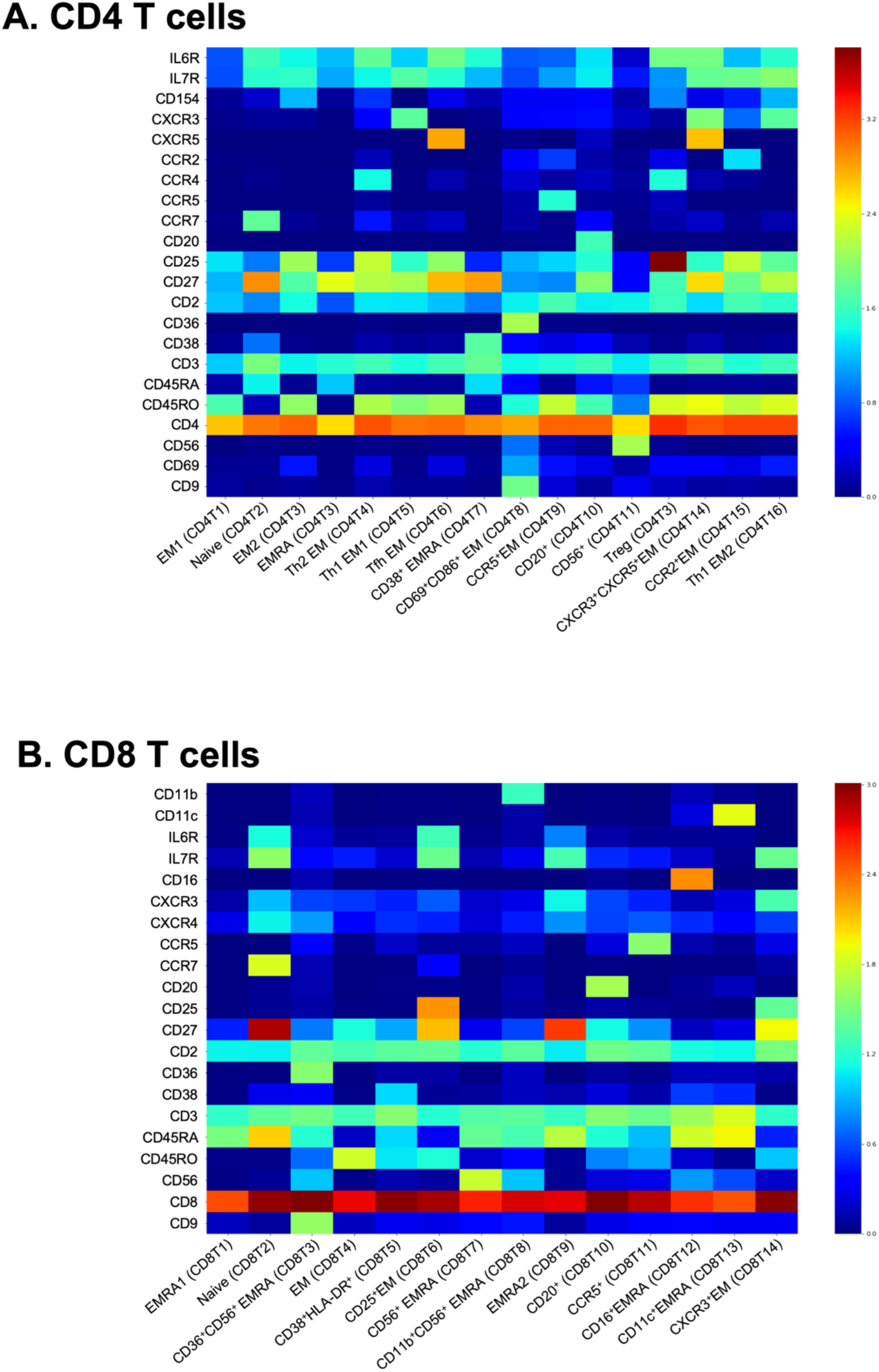
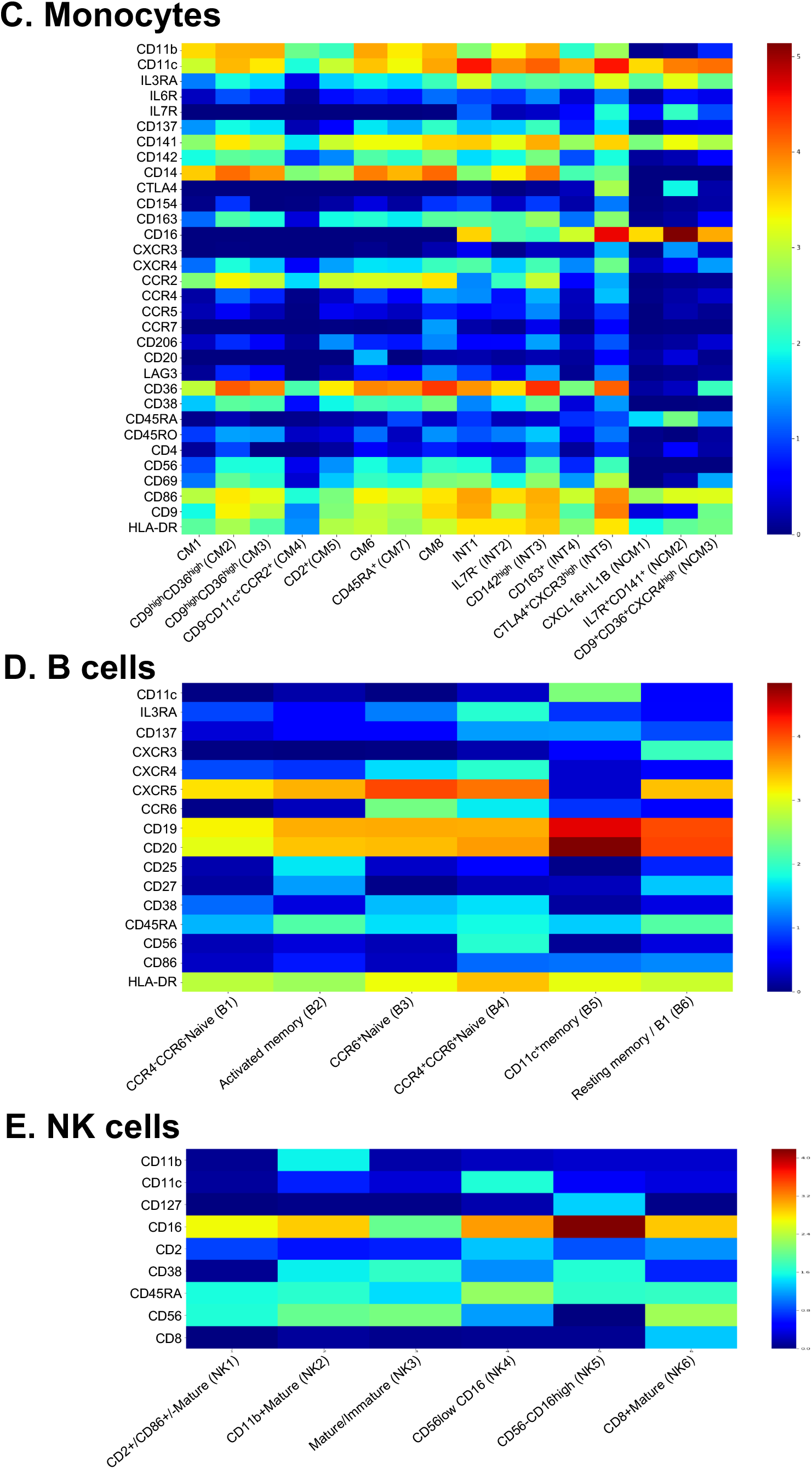
Heatmaps of antibody expression (log_2_ scale) in each main cell type. (**A**) CD4 T cell, (**B**) CD8 T cell, (**C**) Monocytes, (**D**) B cells, and (**E**) NK cells. Immunophenotypes at the bottom. EM, effector memory; EMRA, terminally differentiated effector memory; CM, Classical Monocyte; INT, Intermediate Monocyte; NCM, Nonclassical Monocyte.

Among monocytes, we were able to call 5 of the 8 classical monocyte subsets based on published data.[11] All CM were CD11b+ (**Figure 2C**). There were gradients of CD9, CD69, CD137, CD142 (tissue factor), and CD163 (hemoglobin-haptoglobin receptor) expression. The scavenger receptor CD36, the antigen presentation co-receptor CD86 and the chemokine receptor CCR2 were expressed in all classical monocytes. Based on these markers, 5 of the 8 CM subsets were called (**Figure 2C**) and related to subsets described by mass cytometry. INT CD14+CD16+ monocytes have been considered pro-inflammatory and are known to be increased in people with HIV[12] and with CVD.[41, 51] All INT highly expressed the inflammation-induced costimulatory molecule CD86 (**Figure 2C**). Cluster INT3 highly expressed CD142 (tissue factor), which has previously been implicated in people living with HIV.[45] Since INT subsets have not been described before, we propose a provisional naming suggestion (**Figure 2C**). NCM formed 3 clusters (**Figure 2C**). Strikingly, expression of CD9 and CD36 was limited to cluster 3, suggesting that this cluster corresponds to the previously described CD9+CD36+ NCM.[11] CD11c, CD74, CD86, and CD141 were expressed in all NCMs (**Figure 2C**).

We were able to call all 6 B cell subsets. As expected, CD20 and CD74 (HLA-DR) were expressed in all B cells (**Figure 2D**). CD27, IgM and IgD are used to identify naïve B cells (CD27-IgM+IgD+). Clusters 1, 3, and 4 were negative for CD27 with high transcript expression for IgM and IgD, consistent with naïve B cells. Clusters 3 and 4 expressed CCR6, a subset found in HIV+ subjects.[32] B cell cluster 2 expressed CD25, which is a known marker for B cell proliferation and exhaustion, and CD27, identifying cluster 2 as a likely activated memory B cell. Cluster 5 had high CD11c levels, known to increase in HIV-infected subjects,[19] and expressed some CXCR3 and CCR6, but was CD27low. These features together with moderate expression of CD22 transcript suggest that cluster 5 may contain CD11c+ pathologic B cells. (**Figure 2D**). Most NK cells were mature (CD56^dim^/CD16+), as expected (**Figure 2E**). Cluster 3 also contained immature (CD56^bright^CD16-) NK cells. The CD56^low^CD16− cells (clusters 4 and 5) expressed CD2 and CD45RA. Cluster 5 was CD56-CD16^high^, an NK cell subset known to be elevated in chronic HIV infection.[17] Taken together, this demonstrates the power of combined antibody and transcriptome sequencing.

### Changes in PBMC subset abundance on disease or treatment

Based on this data, it is possible to address shifts in cell proportion based on disease or treatment. We found significant differences in cell proportions in 3 intermediate monocyte subsets, one CD8 T cell, one B cell and one NK cell subset (**Figure 3**). Strikingly, three subsets of intermediate monocytes (**Figure 3A**) showed significantly different abundances. INT2 (IL7R-) and INT3 (TF^hi^) were significantly elevated in WIHS participants living with HIV and drastically reduced in those that also had subclinical CVD. INT4 had an opposite pattern: These CD163− cells were rare in WIHS participants living with HIV, but more abundant in those that also had subclinical CVD. Among B cells, activated memory B2 cells (**Figure 3C**) were severely lower in all WIHS participants with HIV with or without subclinical CVD.

**Figure 3.**
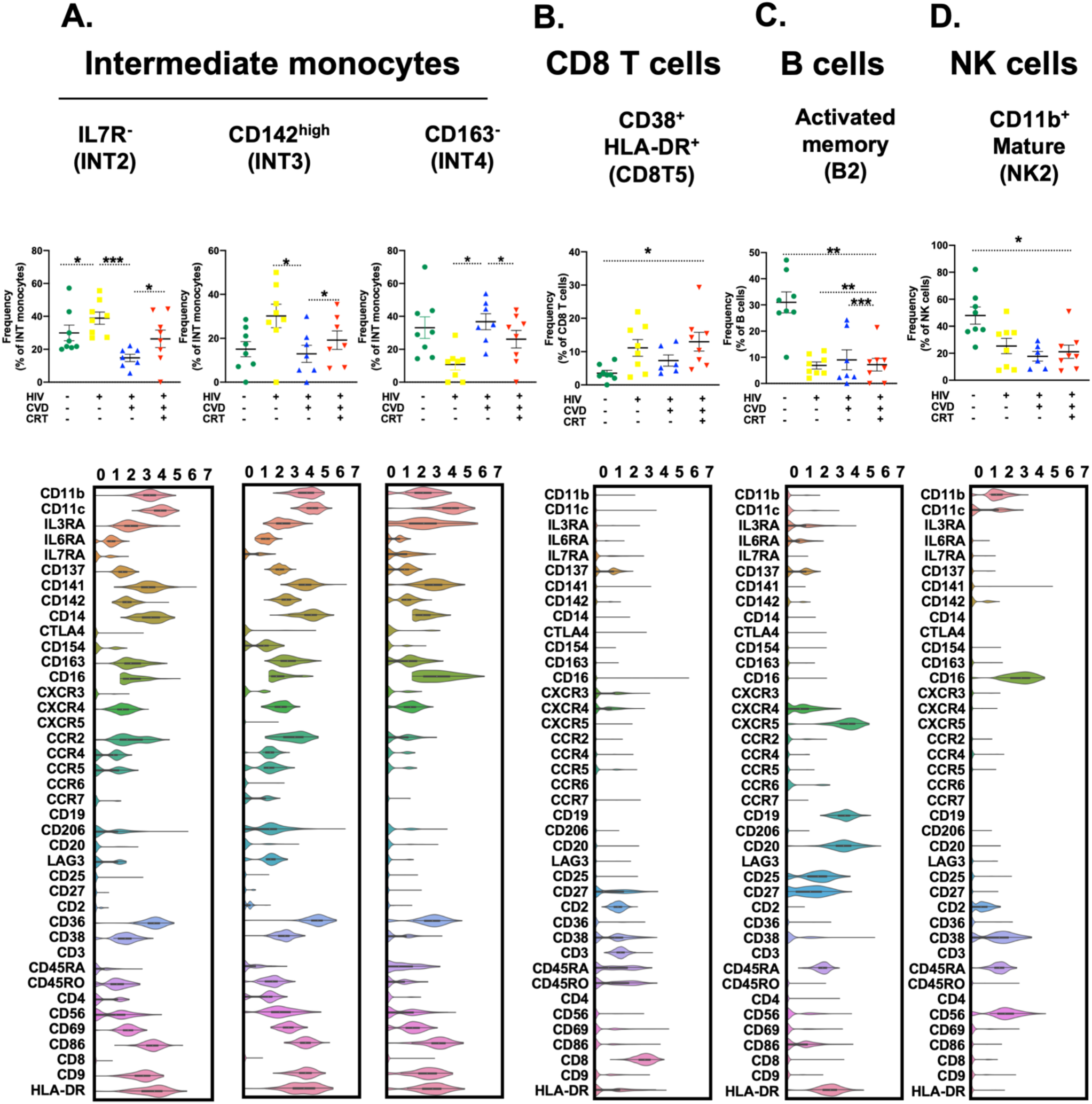
Cell proportions in women with HIV, CVD, both or neither. HIV-CVD− (green), HIV+CVD− (yellow), HIV+CVD+ (blue), and HIV+CVD+ on CRT (red), from left to right. 8 samples per group except 7 for HIV+CVD+. Proportions of cells in each cluster calculated as percentage of the parent cell type as indicated in the title of each panel. Clusters with significant differences (*, p<0.05, **, p<0.01, *** p<0.001) in cell proportions (by log odds ratio) are shown with individual data points, means and standard error of the mean (SEM). Violin plots below show expression of all 40 cell surface markers (log_2_ scale). INT, intermediate monocytes; CRT, cholesterol-reducing treatment.

### Differential gene expression in each of the clusters

Since the transcriptomic information was not used for UMAPs and clustering shown in **Figures 1** and **2**, we were able to compare the gene expression patterns in each cell subset within the same cell type. We filtered for genes that were significantly differentially expressed in at least one of the subsets (**Figure 4, Data S1**). This analysis revealed gene signatures for most subsets. Such gene signatures can then be used to determine the presence of each subset in bulk transcriptomes, and to determine their proportions using Cibersort.[34] As an example, we applied the classical monocyte transcriptomes (8 subsets) to bulk transcriptomes of sorted classical monocytes from 92 subjects.[7] We found 1 of the CM subsets in all subjects and others at varying proportions (**Figure 5**).

**Figure 4.**
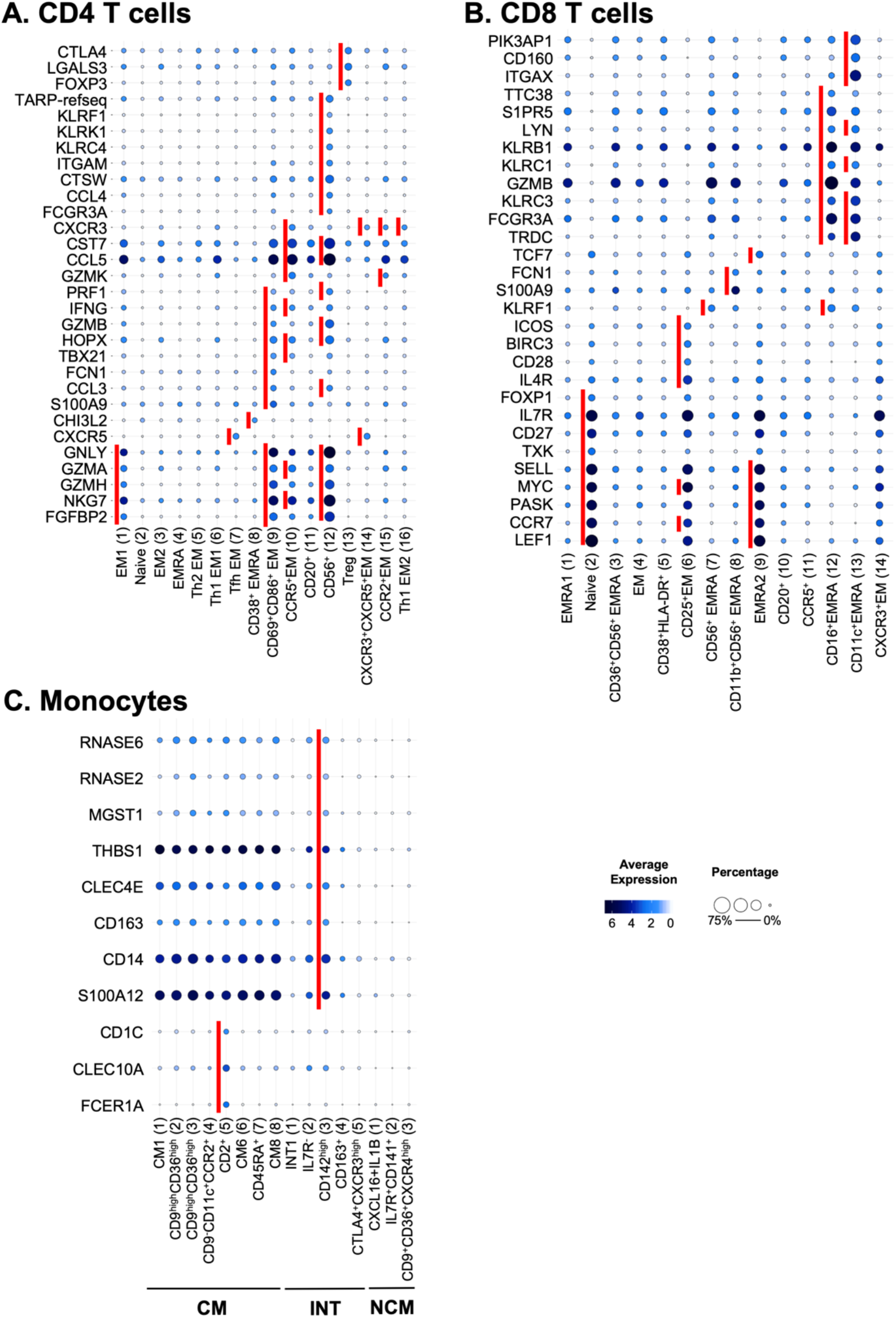
Significantly differentially expressed genes of cells in each cluster. Expression of 485 transcripts was determined by targeted amplification (BD Rhapsody system). Significant genes defined as adjusted p<0.05 and log_2_ fold change >0. Dot plot: fraction of cells in cluster expressing each gene shown by size of circle and level of expression shown from white (=0) to dark blue (=max, log_2_ scale). Red bars indicate genes that were significantly higher in one cluster compared to all other clusters of the parent cell type. There were no DEGs in NK cell clusters. (**A**) CD4 T cells, (**B**) CD8 T cells and (**C**) monocytes. CM, Classical monocytes; INT, Intermediate monocytes; NCM, Nonclassical monocytes; EM, effector memory; EMRA, terminally differentiated effector memory.

**Figure 5.**
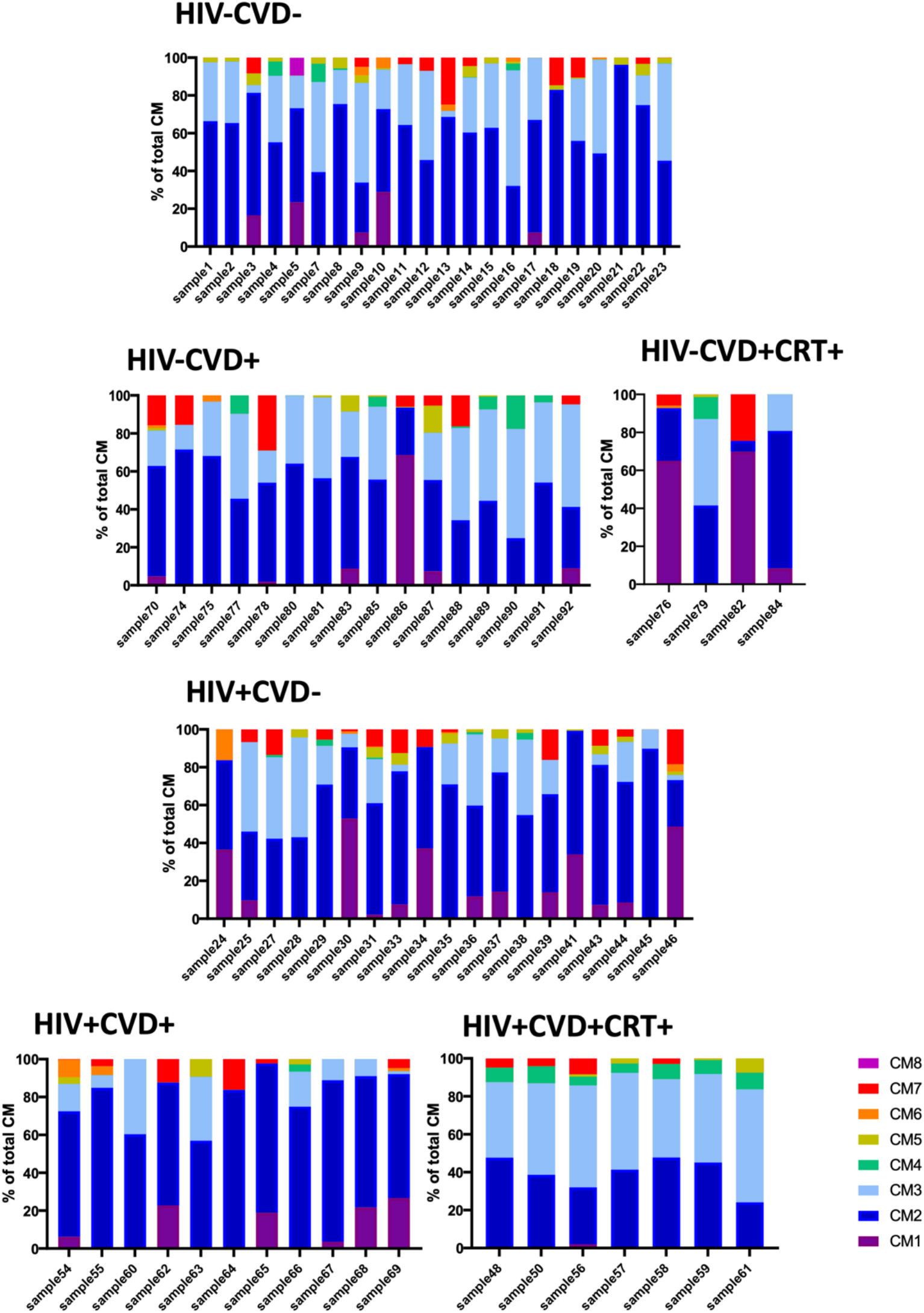
Cell proportions of classical monocyte subsets. Cell proportions of classical monocytes (CMs, CM1-8) from the present dataset in 92 samples of classical monocyte transcriptomes in women with HIV, CVD, both or neither.[7] CM, classical monocyte; CRT, cholesterol-reducing treatment.

### Transcriptomes shift with HIV, CVD and cholesterol control

Transcriptomes may also change with disease state. To test this possibility, we constructed volcano plots (log fold change on the x axis versus -log p on the y axis (**Figure 6, Data S2**). To test for changes with cardiovascular disease (CVD), we plotted genes significantly different between subjects with and without CVD. All these subjects were HIV+. Many genes in CD4 and CD8 T cell subsets showed significant differences. Some genes in monocyte and B cell subsets showed significant differences. To test whether our method could detect effects of treatment, we interrogated transcriptomes of subjects with CVD that received CRT. Again, many genes in T cell and monocyte subsets and some in B cell subsets showed significant differences subjects (**Figure 6**). In CD4T1, 2, and 8, IL-32 was highly significantly increased by CVD, but not in CVD+ women on CRT (**Figure 6**). IL-32 is an inflammatory cytokine that is known to be important in CVD.[6, 21] In CD4T2, L-selectin (SELL), PSGL-1 (SELPLG), and CCR7 were also highly significantly increased in WIHS participants with HIV and CVD. In addition to SELL and SELPLG, CD4T8 showed strong upregulation of TNFSF10 (TRAIL). In CD8T1 and 2, IL32 was high in women with CVD, but less so in women receiving CRT. Other genes highly induced by CVD in CD8T1 included CD52, TRAC and HOPX. Several killer cell lectin receptors (KLRC4, KLRD1, KLRG1 and KLRK1) were also significantly upregulated in CVD. In CD8T3, CD52, CCL5, IL32 and CD160 were all significantly higher in CVD+ participants. CCL5 encodes the chemokine RANTES, known to be important in atherosclerosis.[54] In CD8T4, CVD was associated with significantly increased IL32, TRAC, HOPX, CCL5 and the killer lectin receptors KLRK1, KLRC4, KLRD1.

**Figure 6.**
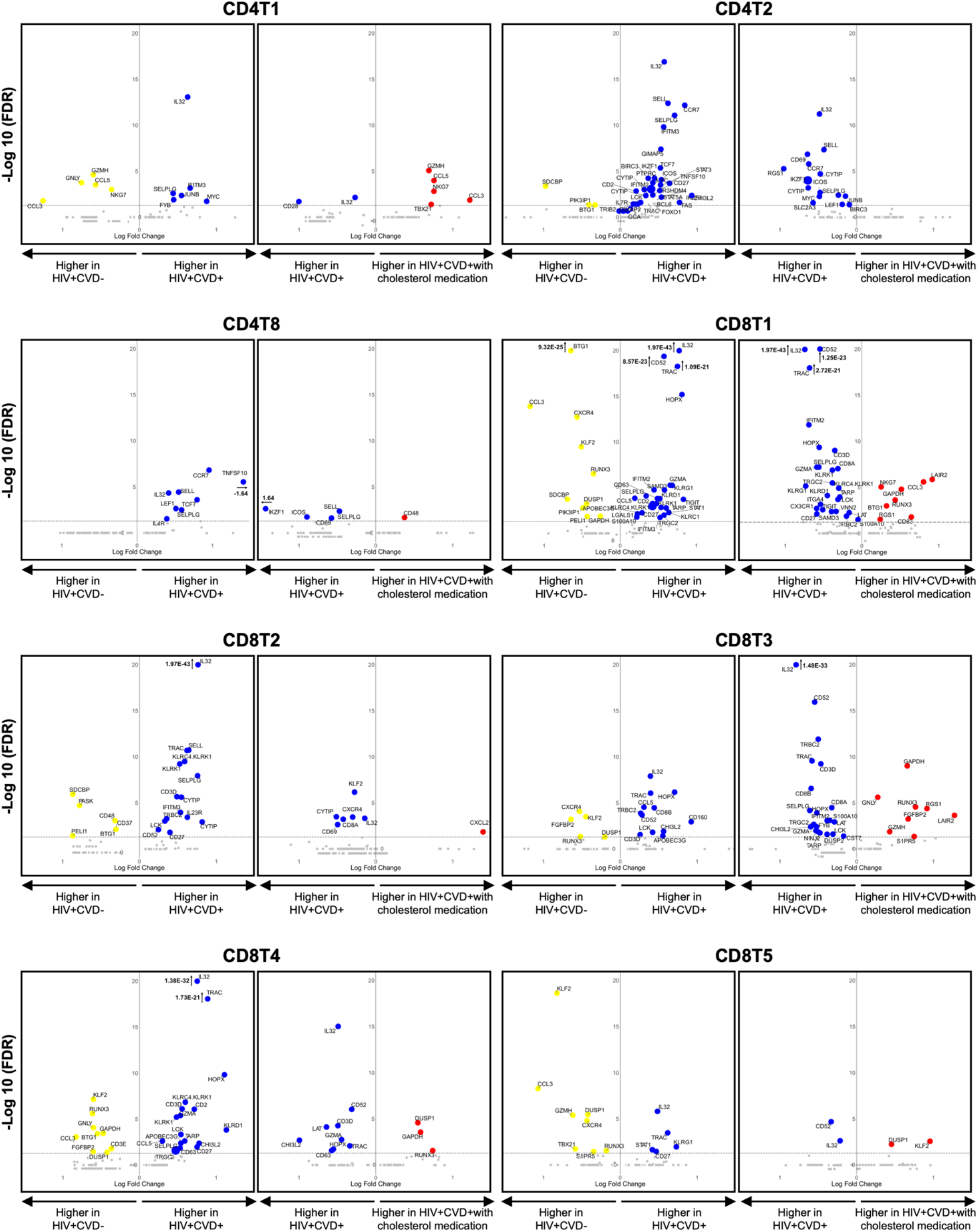
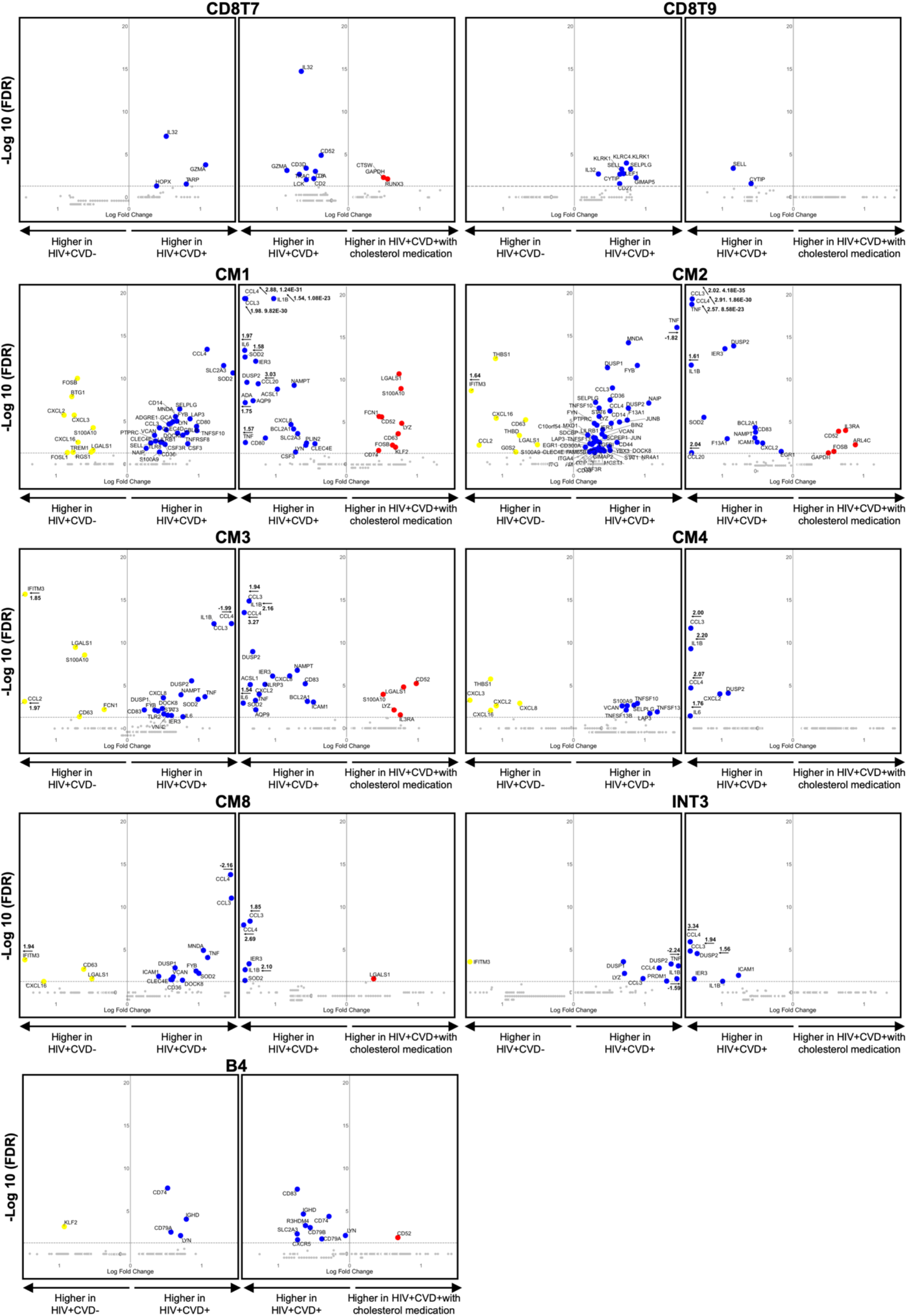
Volcano plots comparing gene expression in single cells from WIHS participant types in each cluster. Gene expression focused on HIV+CVD− vs HIV+CVD+, and HIV+CVD+ vs HIV+CVD+ with CRT. All clusters in which at least 10 genes were statistically significant are shown. Colored dots (HIV+CVD-yellow, HIV+CVD+ blue, and HIV+CVD+ with CRT red) indicate significantly differentiated expressed genes (FDR<0.05 and |log_2_FC|>2). 3 CD4 T and 7 CD8 T cell clusters, 5 CM and 1 each INT and B cell clusters met these criteria. Dashed line indicates adjusted p-value of 0.05. CM, Classical monocytes; INT, Intermediate monocytes.

In CM1, CVD was associated with significantly increased CCL4, SLC2A3, SOD2, and SELPLG. CRT was associated with lower expression of these genes. In CM2, TNF, DUSP1, and 2 were highly associated with CVD (**Figure 6**), as were TNFSF10 (TRAIL), TNFSF13 (APRIL), and TNFSF13B (BAFF), important B cell regulators. In addition to CCL3, CCL4, and DUSP2, IL1B, known to be highly relevant in atherosclerosis, was highly upregulated in CM3 of HIV+CVD+ participants. The Toll-like receptor TLR2, which is known to be involved in atherosclerosis, was upregulated by CVD in CM3. In INT3, CCL3, CCL4, TNF, IL1B, and DUSP2 were associated CVD in the participants that did not receive CRT.

## Discussion

In immunology, surface markers are widely used to define and distinguish cell types.[5, 47, 57] Flow cytometry is the discipline-defining method of immunology.[40] Similar to flow cytometry, in CyTOF, single-cell suspensions are stained with antibody panels to detect cellular antigens. Unlike CyTOF, scRNA-seq allows the detection of single-cell transcriptomes. Since the correlation between cell surface protein and mRNA expression is weak in immune cells[25], the transcriptome provides a valuable additional dimension. scRNA-Seq without surface phenotype information has led to much frustration in the field, because the expression of many genes encoding well-known surface markers remains undetected in scRNA-Seq.[27, 57] It is still difficult to call cell types based on gene expression data alone, which emphasizes the need for cell surface phenotypes in addition to transcriptomes. Here, we correlated gene expression with cell surface expression for 41 pairs of genes and proteins. CD74 surface expression was well correlated with the expression of both the *CD74* and the *HLA-DRA* genes. CD4 and CD16 surface and gene expression were reasonably well correlated across all cell types. A few other genes including *CD14*, *CD16*, IL-3 receptor (*CD123*), and *CD27* were somewhat correlated with the surface expression of their proteins in some cell types. For most markers, we confirm weak correlations,[25] which illustrates the value of monitoring cell surface phenotype in scRNA-Seq.

PBMCs can be analyzed without mechanical or enzymatic dissociation, which are known to alter cell surface markers and transcriptomes.[56] PBMC are attractive for single cell RNA sequencing (scRNA-Seq) studies, because they are available in many clinical studies of specific populations with defined diseases and outcomes. The participants sampled for the present study were part of a sub-study nested within the WIHS,[13, 16, 18] which provided detailed information on subclinical atherosclerosis. Participants underwent high-resolution B-mode carotid artery ultrasound to image six locations in the right carotid artery.[16] Although our study is not definitive, it is suggestive of significant changes in cell proportions and transcriptomes in subjects with cardiovascular disease.

scRNA-Seq has been applied to human PBMCs in diseases including cancers,[3, 60–62] inflammatory bowel disease,[30, 53] and autoimmune disease,[22, 38] as well as atherosclerosis.[9, 58] The foundational paper for the 10x Genomics drop-Seq method[63] demonstrated the feasibility of using scRNA-Seq on PBMCs. Other studies reported scPBMC transcriptomes in colorectal cancer,[61] γδ T cells,[36] liver cancer,[62] in vitro salmonella infection,[2] and memory T cells.[28] Only two publications reported single cell transcriptomes from patients with atherosclerosis (carotid endarterectomy specimens and matched PBMCs).[9, 58] 1,652 PBMCs from one patient were analyzed by 10x Genomics 3’ and cellular indexing of transcriptional epitope sequencing (CITE-Seq),[35, 48] using a panel of 21 antibodies. No healthy control PBMCs were studied. ScRNA-Seq revealed that the process of smooth muscle cell phenotypic modulation *in vivo* can be altered by the expression of *Tcf21*, a gene causally associated with reduced risk of coronary artery disease. The loss of *Tcf21* results in fewer fibromyocytes in the lesions and the protective fibrous cap.[58] A recent study reported the effect of HIV infection on PBMC transcriptomes,[20] focusing on acute HIV infection (before antiretroviral therapy started) and reporting PBMC transcriptomes in four patients at 8 defined time points (average of 1,976 PBMC transcriptomes per participant and condition). No scRNA-Seq or CITE-Seq studies of PBMCs of people living with chronic HIV infection have been reported. No single cell studies of the interaction between HIV and CVD are available.

Six clusters showed significantly different abundance of cells in the four groups of participants, three of them intermediate monocyte subsets, which underscores the extraordinary importance of this cell type in chronic HIV infection[14, 29] and CVD.[10, 24] Intermediate monocyte numbers have previously been found increased in non-HIV subjects with peripheral artery occlusive disease[55] and significantly predicted cardiovascular events.[15, 41, 42] Cells in INT1, the largest cluster, shared CD11b, CD11c, CD9, CD36, CD38, CD56, CD69, CD83, IL-3RA, IL6R, CD137, CD141, CD142 (tissue factor), CXCR4 and CD74 (HLA-DR) with other intermediate monocytes. We found no single positive marker that was specific for INT1 and thus refrained from naming this cluster. We found the INT2 and INT3 increased in women living with HIV. Both express tissue factor (CD142). Tissue factor expression on monocytes has previously been shown to be increased in HIV+ subjects.[45] Intermediate monocytes are considered pro-atherogenic,[44] and tissue factor expression provides a plausible reason for this. We found that in INT2 and INT3, the inflammatory chemokines CCL3 and CCL4 and the known pro-atherogenic cytokine IL-1β were significantly upregulated in participants with CVD, but not in those receiving CRT. INT4 uniquely lack expression of CD163, the receptor for hemoglobin-haptoglobin complexes. Thus, we call INT4 CD163-intermediate monocytes. INT5, called (CTLA4+CXCR3hi) uniquely expressed CTLA4 (CD152) and highly expressed CXCR3.

In CD4T cells clusters 1, 2 and 8, IL-32 was highly significantly increased by CVD, which was reversed by cholesterol lowering in CD4T1 and 2 (**Figure 6**). IL-32 is a 27 kDa cytokine expressed in T cells and monocytes that is secreted after apoptosis.[33] It is an inflammatory cytokine that drives IL-1β, clinically important in CVD,[39] TNF, IL-6 and IL-8 expression.[6, 21, 33] IL-32 activates the leukocyte surface protease PR3, which in turn triggers the G-protein coupled receptor PAR2[33] and is known to be important in viral infections.[23, 33, 37, 46] We found IL-32 highly expressed in most T and NK cell clusters. Since IL-32 appears to be CVD-specific, we advocate for future prospective studies in larger cohorts to determine whether IL32 mRNA is a useful biomarker.

Our discovery study will encourage prospective epidemiological studies to address which PBMC subset and transcriptomes are best suited as clinical biomarkers to stratify risk and guide treatment in subjects with coronary or peripheral artery disease. The current findings also present some limitations. They need to be extended to men (the current data is based on women) and other races and ethnicities (the current data is based on mostly African American and Hispanic women). Studies of CVD in non-smokers are also needed (the current data is based on smokers), and the age range needs to be broadened.

In conclusion, we demonstrate the utility of scRNA-Seq with cell surface phenotype assessment in the same cells. The identification of 58 distinct clusters of CD4 and CD8 T cells, B cells, NK cells and monocytes helps to gain a deeper understanding of PBMCs, a rich and readily accessible source of biological and clinical information. The discovery of subsets of intermediate monocytes calls for identifying such subsets in model organisms to test their function *in vivo*.

## Declarations

### Funding

National Institutes of Health Grant R35-HL-145241, R01-HL-121697, R01-HL-148094 (K.L.)

National Institutes of Health Grant P01-KL-136275 (C.C.H.)

National Institutes of Health Grant R01-HL-134236 (C.C.H.)

National Institutes of Health Grant R01-HL-126543, 5R01-HL-126543-05, 5R01-HL-140976-02, R01-HL-148094-01, R01-HL-148094 (R.C.K.)

National Institutes of Health Grant K01-HL-137557 (D.B.H.)

National Institutes of Health Grant U01-AI-103408 (I.O.)

National Institutes of Health Grant U01-AI-103390 (A.A.A.)

National Institutes of Health Grant U01-AI-034989 (P.C.T.)

Cancer Research Institute (CRI) (A.S.)

American Heart Association (AHA) 19POST34450020 (L.E.P)

National Institutes of Health Grant NIAID, NICHD, NCI, NIDA, NIMH, NIDCR, NIAAA, NIDCD, UL1-TR-000004, P30-AI-050409, P30-AI-050410, and P30-AI-027767.

Formación de Profesorado Universitario (FPU) 16/02780 (R.B.D.)

Swedish Society for Medical Research (SSMF) (J.V.)

Data in this manuscript were collected by the Women’s Interagency HIV study, now the MACS/WIHS Combined Cohort Study (MWCCS).

### Author contributions

Design of the study: J.V., R.S., Y.G., C.P.D., E.E.

Collection of samples and data. A.L.L., R.P.T., J.M.L., W.J.M., K.M.W, A.A.A., H.N.H., P.C.T., I.O., S.L.H., and R.C.K.

Analysis of clinical data: D.B.H.

Design and collection of data for the B mode ultrasound sub-study: H.N.H. scRNA-Seq experiments: C.P.D., and E.E.

Analysis of the data: J.V., R.S., R.G., Y.G., P.R., T.P., L.E.P., C.E.O., R.B.D., H.Q.D., A.S., C.A.M., L.L.L., T.W., C.C.H., and K.L.

Bioinformatics analysis: R.G., Y.G. and H.Q.D.

Writing-original draft: J.V., R.S., and K.L.

### Data and material availability

All data are available in the main text or the supplementary materials.

### Code availability

All packages used for the analysis on this data are available in R.

### Ethics approval

All participants provided informed consent, and each site’s Institutional Review Board approved the studies.

### Consent to participate

All participants provided informed consent, and each site’s Institutional Review Board approved the studies.

### Consent for publication

All the listed authors have reviewed the manuscript and agreed with its submission.

