## Supplemental material (figures and tables) for "Combined protein and transcript single cell RNA sequencing in human peripheral blood mononuclear cells"

**Long title**

Klaus Ley, MD

La Jolla Institute for Immunology

9420 Athena Circle

La Jolla, CA 92037, USA

(858) 752-6661 (tel)

(858) 752-6985 (fax)

**This PDF file includes:**

Figs. S1 to S3
Tables S1 to S11

**Other Supplementary Materials for this manuscript include the following:**

Data S1 to S2

**Fig. S1.**

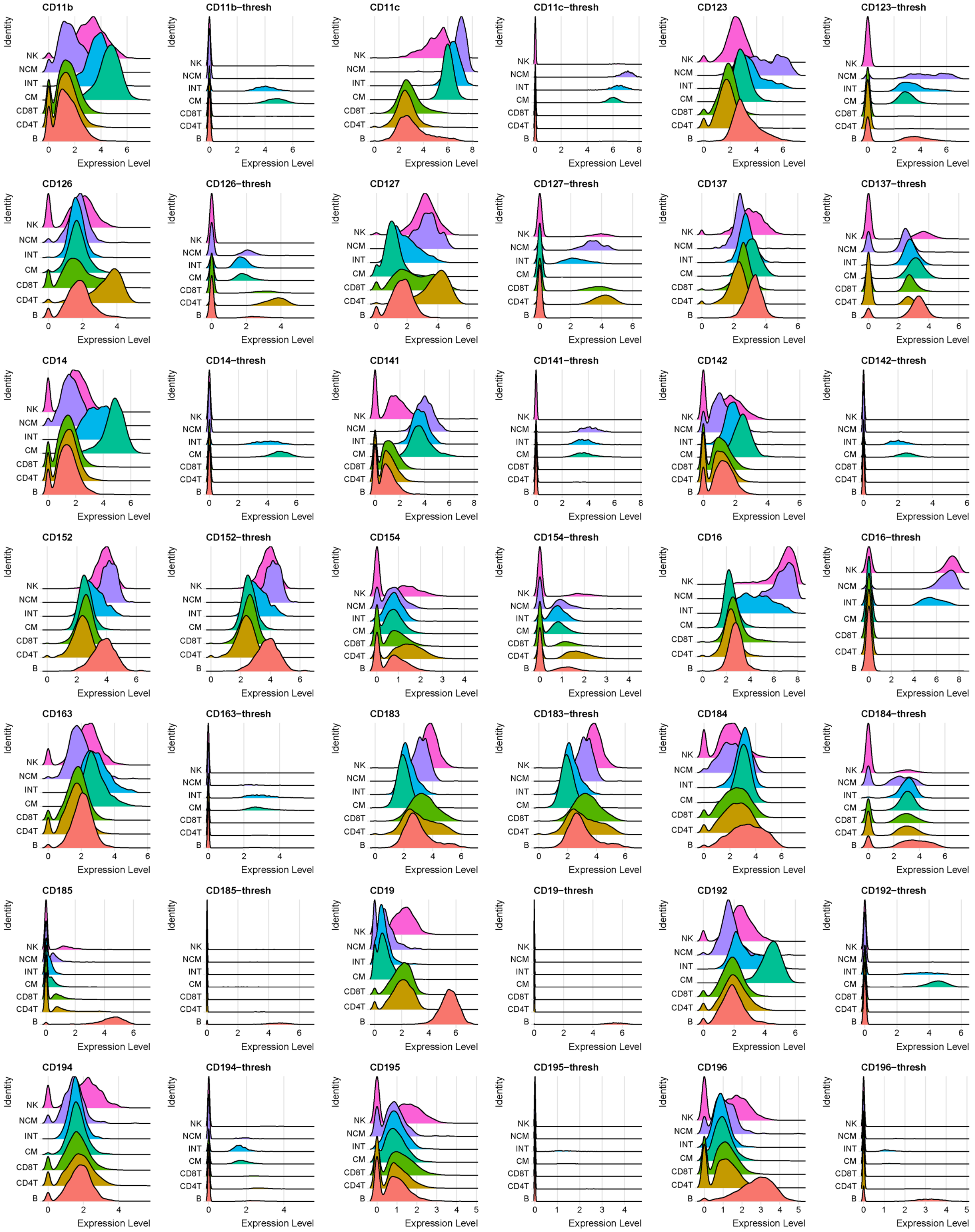
CD11b
CD11b−thresh
CD11c
CD11c−thresh
CD123
CD123−thresh
CD126
CD126−thresh
CD127
CD127−thresh
CD137
CD137−thresh
CD14
CD14−thresh
CD141
CD141−thresh
CD142
CD142−thresh
CD152
CD152−thresh
CD154
CD154−thresh
CD16
CD16−thresh
CD163
CD163−thresh
CD183
CD183−thresh
CD184
CD184−thresh
CD185
CD185−thresh
CD19
CD19−thresh
CD192
CD192−thresh
CD194
CD194−thresh
CD195
CD195−thresh
CD196
CD196−thresh
Identity
NK
NCM
INT
CM
CD8T
CD4T
B
Expression Level

### Fig. S1 cont.

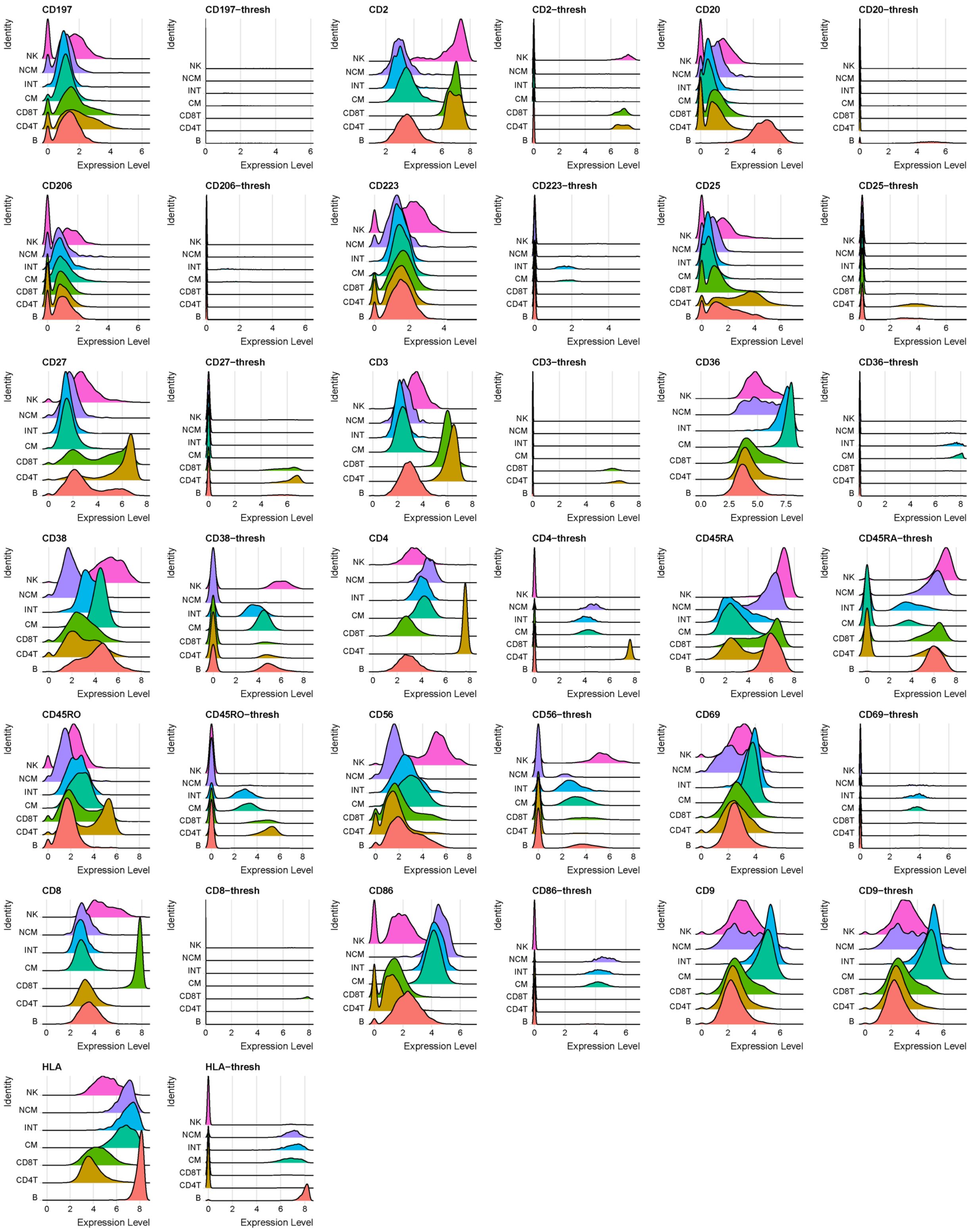
CD197
CD197−thresh
CD2
CD2−thresh
CD20
CD20−thresh
CD206
CD206−thresh
CD223
CD223−thresh
CD25
CD25−thresh
CD27
CD27−thresh
CD3
CD3−thresh
CD36
CD36−thresh
CD38
CD38−thresh
CD4
CD4−thresh
CD45RA
CD45RA−thresh
CD45RO
CD45RO−thresh
CD56
CD56−thresh
CD69
CD69−thresh
CD8
CD8−thresh
CD86
CD86−thresh
CD9
CD9−thresh
HLA
HLA−thresh
Identity
NK
NCM
INT
CM
CD8T
CD4T
B
Expression Level

**Figure S1. Ridgeline plots of the unthresholded and thresholded expressions of all the 40 antibodies for each main cell types** [CD4 T cells, CD8 T cells, classical monocytes (CM), intermediate monocytes (INT), nonclassical monocytes (NCM), NK cells and B cells] using Seurat. These plots separately show the distribution of each value of CLR normalized antibody derived tag for each main cell type. Expression levels are shown on x-axis. Thresh, thresholded.

**Fig. S2.**

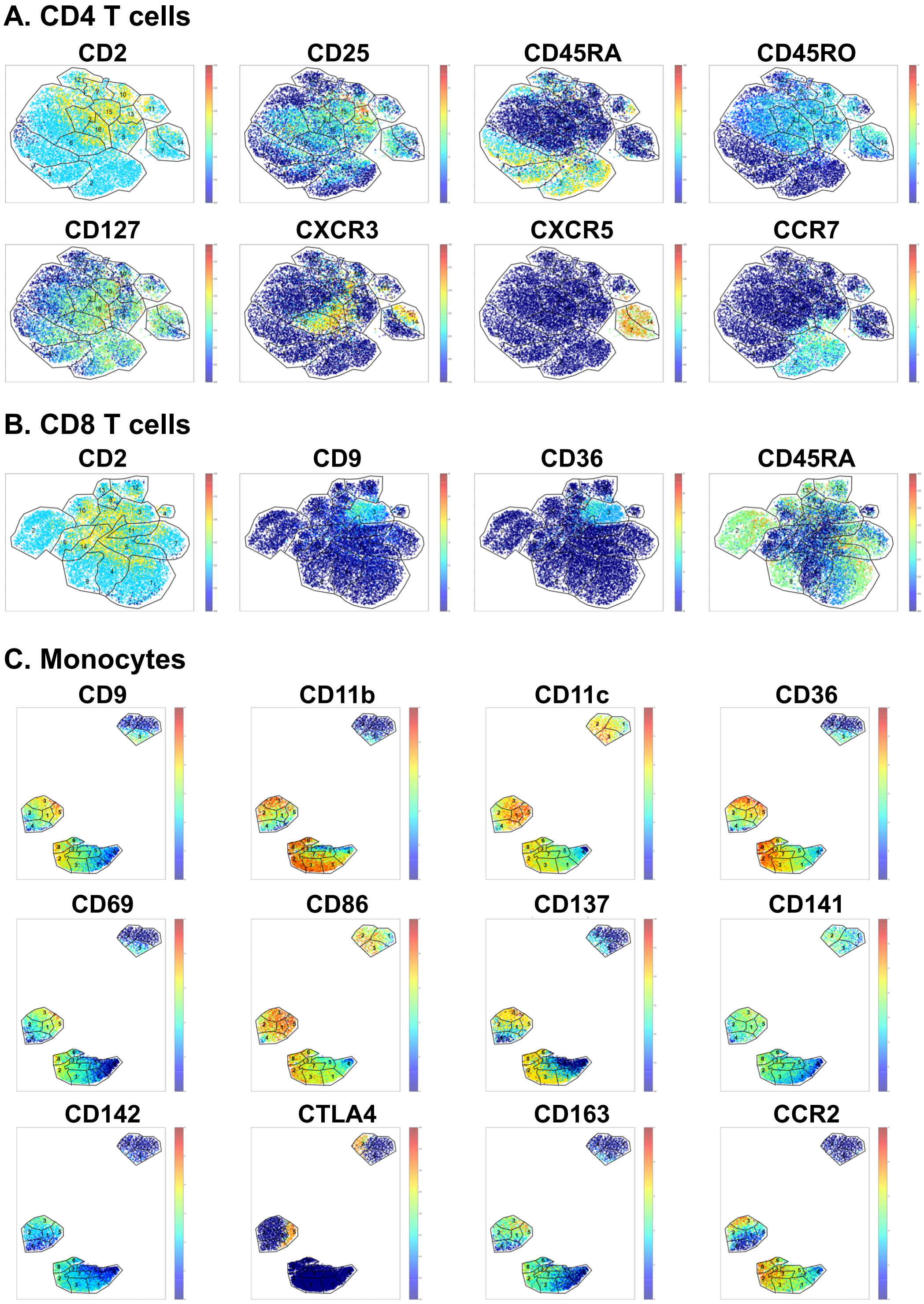
A. CD4 T cells
CD2
CD25
CD45RA
CD45RO
CD127
CXCR3
CXCR5
CCR7
B. CD8 T cells
CD2
CD9
CD36
CD45RA
C. Monocytes
CD9
CD11b
CD11c
CD36
CD69
CD86
CD137
CD141
CD142
CTLA4
CD163
CCR2

**Fig. S2 cont.**

#### D. B cells

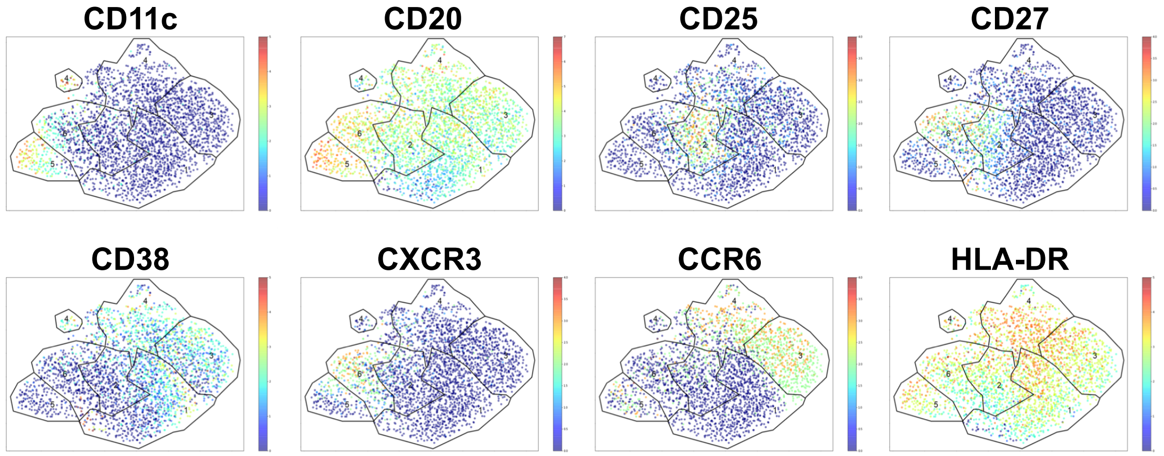


#### E. NK cells

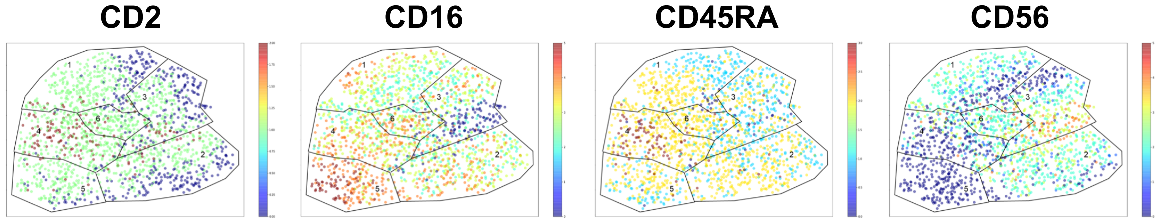


**Fig S2. Cell surface marker expression.** The expression level of each of the 40 antibody markers was color-coded from dark blue (=0, not expressed) to red (highest expression, log2 scale, as per color bar in each panel). Feature plots projected on UMAP gates of each cell type. Selected surface markers shown on top of each plot. Cluster outlines as defined in Figure 2. (**A**) CD4 T cells; (**B**) CD8 T cells; (**C**) Monocytes; (**D**) B cells, and (**E**) NK cells.

**Fig. S3.**

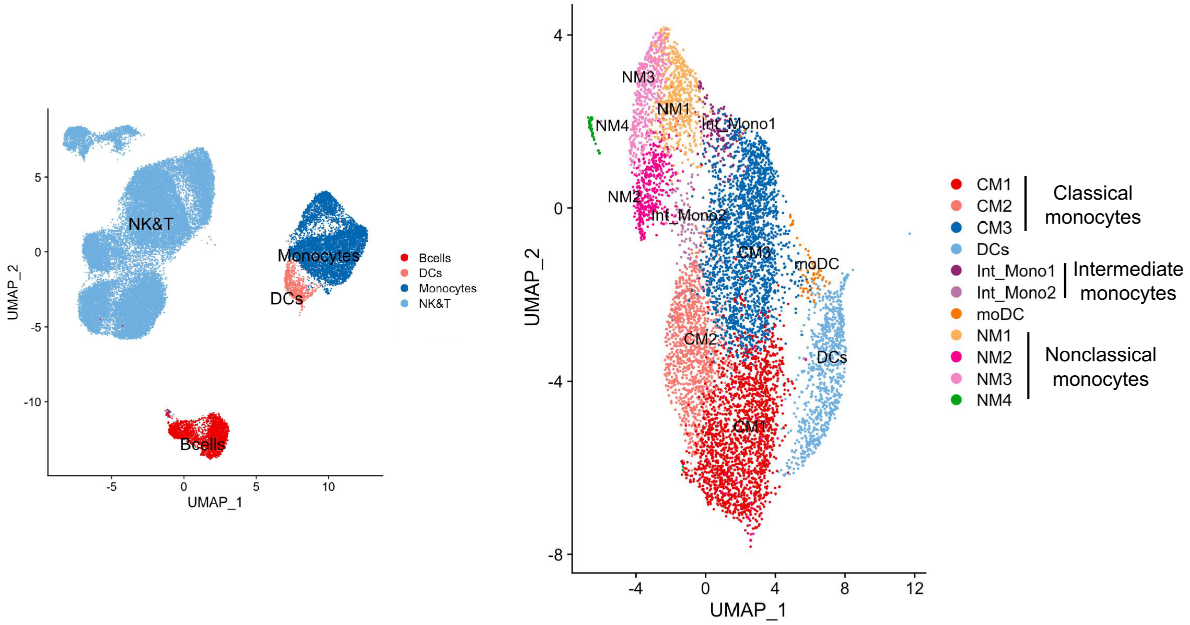


**Fig S3. Unbiased UMAP clustering.** UMAP clustering based on 462 genes and 40 antibody markers using Seurat, revealing B cells, monocytes, DCs, NK cells and T cells (left). The monocytes and DCs were re-clustered showing 3 classical, 2 intermediate and 4 nonclassical monocyte clusters, 1 DC and 1 monocyte-DC cluster (right). CM, Classical monocytes; INT, Intermediate monocytes; NCM, nonclassical monocytes; DCs (Dendritic cells).

**Table S1.**

| Reagent | Vendor | Catalogue # |
|---|---|---|
| cRPMI*1 | | - |
| PBS | Fisher Scientific | 10010049 |
| FBS | Gemini | 100106-500mL |
| Trypan Blue | Thermo Fisher | 15250061 |
| Fc Block | BD | 564220 |
| SMK | BD | 633781 |
| DRAQ7 | BD | 564904 |
| Calcein AM*2 | Thermo Fisher | C1430 |
| DMSO | Thermo Fisher | D12345 |
| AbSeq | BD | - |
| Rhapsody Reagent Kit | | 633771 |
| Rhapsodu Human Immune Response Panel | | 633750 |
| Rhapsody Custom Reagent Panel | | 633742 |
| AMPure XP beads | Beckman Coulter | A63881 |
| Ethanol | Milipore Sigma | E7023-500ml |
| Nuclease Free Water | Qiagen | 129114 |
| D1000 ScreenTape | Agilent | 5067-5584 |
| D1000 Sample Buffer | Agilent | 5067-5603 |
| Qubit Reagent Kit | Thermo Fisher | Q33231 |
| NovaSeq S1 100 Cycle Kit | Illumina | 20012865 |
| NovaSeq S2 100 Cycle Kit | Illumina | 20012862 |

**Table S1. Reagents.** *1: From 500mL of RPMI-1640 (with L-Glutamine) remove 75.5 mL and transfer to two 50 mL conical tubes for future potential use. Add 50 mL of 100% human serum albumin (HSA). Add 5 mL of the following (concentrations indicated are of stock solutions): HEPES (1M), sodium pyruvate (100X) MEM-NEAA (100X), penicillin-streptomycin, and GlutaMAX. Add 0.5 mL of mercaptoethanol (1000X). Mix by inversion in RPMI-1640 original bottle. Carefully transfer solution to a 500 mL CorningStore at 4° C. Vacuum filter system with a 0.45 mm filter size. *2: Resuspended in DMSO.

**Table S2.**

| Day 1 | Viability (%) | Day 2 | Viability (%) |
|---|---|---|---|
| 1 | 94 | 1 | 91 |
| 2 | 93 | 2 | 93 |
| 3 | 81 | 3 | 85 |
| 4 | 87 | 4 | 91 |
| 5 | 81 | 5 | 92 |
| 6 | 87 | 6 | 88 |
| 7 | 89 | 7 | 83 |
| 8 | 89 | 8 | 84 |
| 9 | 95 | 9 | 94 |
| 10 | 94 | 10 | 91 |
| 11 | 75 | 11 | 86 |
| 12 | 89 | 12 | 89 |
| 13 | 79 | 13 | 89 |
| 14 | 92 | 14 | 94 |
| 15 | 85 | 15 | 87 |
| 16 | 75 | 16 | 93 |
| **Min** | **75** | **Min** | **83** |
| **Max** | **95** | **Max** | **94** |
| **Median** | **88** | **Median** | **90** |
| **Average** | **87** | **Average** | **89** |

**Table S2: The cell viability of each frozen PBMC tube.** On two separate days 16 tubes of frozen PBMCs were thawed and used. The viability was determined using the BD Rhapsody Scanner. Minimum (Min), maximum (Max), median, and average viability were calculated in each day.

**Table S3.**

| Specificity | Clone | Catalogue Number |
|---|---|---|
| CD11b | M1/70 | 940008 |
| CD11c | B-LY6 | 940024 |
| CD123 (IL-3RA) | 7G3 | 940020 |
| CD126 (IL-6R) | M5 | 940090 |
| CD127 (IL-7R) | HIL-7R-M21 | 940012 |
| CD137 | 4B4-1 | 940055 |
| CD14 | MPHIP9 | 940005 |
| CD141 | 1A4 | 940079 |
| CD142 | HTF-1 | 940280 |
| CD152 (CTLA-4) | BNI3 | 940034 |
| CD154 | TRAP1 | 940053 |
| CD16 | 3G8 | 940006 |
| CD163 | GHI/61 | 940058 |
| CD183 (CXCR3) | 1C6/CXCR3 | 940030 |
| CD184 (CXCR4) | 12G5 | 940056 |
| CD185 (CXCR5) | RF8B2 | 940042 |
| CD19 | SJ25C1 | 940004 |
| CD192 (CCR2) | 1D9 | 940286 |
| CD194 (CCR4) | 1G1 | 940047 |
| CD195 (CCR5) | 2D7/CCR5 | 940050 |
| CD196 (CCR6) | 11A9 | 940033 |
| CD197 (CCR7) | 3D12 | 940014 |
| CD2 | RPA-2.10 | 940046 |
| CD20 | 2H7 | 940016 |
| CD206 | 19.2 | 940068 |
| CD223 (LAG-3) | T47-530 | 940080 |
| CD25 | 2A3 | 940009 |
| CD27 | M-T271 | 940018 |
| CD3 | SK7 | 940000 |

| | | |
|---|---|---|
| CD36 | CB38 (NL07) | 940224 |
| CD38 | HIT2 | 940013 |
| CD4 | SK3 | 940001 |
| CD45RA | HI100 | 940011 |
| CD45RO | UCHL1 | 940022 |
| CD56 | NCAM16.2 | 940007 |
| CD69 | FN50 | 940019 |
| CD8 | RPA-T8 | 940003 |
| CD86 | 2331(FUN-1) | 940025 |
| CD9 | M-L13 | 940078 |
| HLA-DR (CD74) | G46-6 | 940010 |

**Table S3. List of 40 titrated oligonucleotide-tagged monoclonal antibodies.** All the monoclonal antibodies were from BD Biosciences (NJ, USA). A cocktail of these 40 antibodies was used for staining of the cells following manufacturer's recommendations.

**Table S4.**

| Genes |
| --- |
| ACSL1 |
| AL137655 |
| APLP2 |
| APOBEC3A |
| ASAH1 |
| B3GALT2 |
| BC013828 |
| C3AR1 |
| C5AR1 |
| CAMKK2 |
| CAVIN2 |
| CD300C |
| CD43 |
| CD83 |
| CD96 |
| CSF3R |
| CTSA |
| CYTIP |
| EMR1 |
| FBP1 |
| FCGR2A |
| FCGR3B |
| FGFBP2 |
| FPR1 |
| G0S2 |
| GCA |
| GLUL |
| GNG11 |
| GPR56 |

| |
|---|
| HOPX |
| IFITM1 |
| IKZF3 |
| ITGA2B |
| ITGB3 |
| ITM2B |
| JMJD1C-AS1 |
| KLF2 |
| KLRC4-KLRK1 |
| KLRD1 |
| LILRB1 |
| LRP1 |
| LYZ |
| MCTP2 |
| MEG8 |
| MIR4718 |
| MIR5192 |
| MMP25 |
| MMRN1 |
| MNDA |
| MXD1 |
| NAIP |
| NCF1 |
| NCF1C |
| NEAT1 |
| NLRP3 |
| NR4A1 |
| PELI1 |
| PLIN2 |
| PTAFR |
| PXN |

| |
|---|
| R3HDM4 |
| RAB27B |
| RASGEF1B |
| S100A8 |
| S1PR5 |
| SAMD3 |
| SCPEP1 |
| SDCBP |
| SH2D1B |
| SH3BGRL2 |
| SIGLEC10 |
| SLC11A1 |
| SLC2A3 |
| SLC35G2 |
| SOD2 |
| SORL1 |
| SPON2 |
| SRGN |
| SYNE1 |
| SYNE2 |
| TC2N |
| TCRBV3S1 |
| TKT |
| TNFRSF10C |
| TRGC2 |
| TSPAN14 |
| TTC38 |
| TTYH3 |
| VCAN |
| XPO6 |

| |
|---|
| XYLT1 |

**Table S4: List of selected genes included in the custom panel.** A total of 91 genes were included in the customized panel in addition to the already described BD Human Immune Response Panel.

**Table S5.**

| Antibodies | Threshold |
|---|---|
| CD11b | 1.1 |
| CD11c | 1.5 |
| CD123 (IL-3RA) | 1 |
| CD126 | 0.6 |
| CD127 (IL-7R) | 0.8 |
| CD137 | 1 |
| CD14 | 1.5 |
| CD141 | 1 |
| CD142 | 0.8 |
| CD152 (CTLA-4) | 2.4 |
| CD154 | 1.05 |
| CD16 | 1.5 |
| CD163 | 1.1 |
| CD183 (CXCR3) | 1.1 |
| CD184 (CXCR4) | 0.6 |
| CD185 (CXCR5) | 2 |
| CD19 | 2 |
| CD192 (CCR2) | 1 |
| CD194 (CCR4) | 1.1 |
| CD195 (CCR5) | 1.1 |
| CD196 (CCR6) | 1.9 |
| CD197 (CCR7) | 1.3 |
| CD2 | 0.3 |
| CD20 | 1.25 |
| CD206 | 1.25 |
| CD223 (LAG-3) | 1.25 |
| CD25 | 1 |
| CD27 | 0.5 |
| CD3 | 0.5 |

| | |
|---|---|
| CD36 | 1.5 |
| CD38 | 1.1 |
| CD4 | 1.3 |
| CD45RA | 0.5 |
| CD45RO | 0.75 |
| CD56 | 1.1 |
| CD69 | 1 |
| CD8 | 0.9 |
| CD86 | 0.9 |
| CD9 | 0.8 |
| CD74 (HLA-DR) | 1.2 |

**Table S5. Threshold values of each antibody.** For the antibodies we use CLR Normalization, which is the ratio of the UMI value to the geometric mean of all the UMIs in that cell in log scale. The thresholds are in the same space.

**Table S6.**

| B | CD4 T | CD8 T | CM | INT | NCM | NK |
|---|---|---|---|---|---|---|
| CD3 | CD8 | CD4 | CD3 | CD3 | CD3 | CD3 |
| | CD19 | CD19 | CD16 | CD19 | CD14 | CD14 |
| | | | CD19 | | CD19 | CD19 |
| | | | | | CD56 | CD152 (CTLA-4) |
| | | | | | | CD196 (CCR6) |

**Table S6. Antibodies not used for cell clustering.** B; B cells (CD19+CD3-), CD4 T; CD4 T cells (CD19-CD3+CD4+CD8-), CD8 T; CD8 T cells (CD19-CD3+CD4-CD8+), CM; classical monocytes (CD3-CD19-CD14+CD16-), INT; intermediate monocytes (CD3-CD19-CD14+CD16+), NCM; nonclassical monocytes (CD3-CD19-CD56-CD16+), NK; NK cells (CD56+CD14-CD20-CD123-CD206-).

**Table S7.**

| | B cells | CD4 T cells | CD8 T cells | Classical Monocytes | Intermediate monocytes | Nonclassical monocytes | NK cells |
|---|---|---|---|---|---|---|---|
| **Cluster1** | 884 | 2065 | 2387 | 1310 | 139 | 176 | 446 |
| **Cluster2** | 370 | 1732 | 1555 | 1075 | 258 | 165 | 439 |
| **Cluster3** | 635 | 641 | 1416 | 748 | 242 | 134 | 330 |
| **Cluster4** | 526 | 1074 | 1413 | 600 | 221 | | 247 |
| **Cluster5** | 178 | 792 | 1361 | 498 | 149 | | 212 |
| **Cluster6** | 326 | 913 | 660 | 158 | | | 169 |
| **Cluster7** | | 470 | 943 | 431 | | | |
| **Cluster8** | | 723 | 198 | 325 | | | |
| **Cluster9** | | 269 | 737 | | | | |
| **Cluster10** | | 286 | 611 | | | | |
| **Cluster11** | | 272 | 507 | | | | |
| **Cluster12** | | 329 | 480 | | | | |
| **Cluster13** | | 211 | 142 | | | | |
| **Cluster14** | | 336 | 433 | | | | |
| **Cluster15** | | 478 | | | | | |
| **Cluster16** | | 454 | | | | | |
| **Total** | 2919 | 11045 | 12843 | 5145 | 1009 | 475 | 1843 |

**Table S7: Number of Cells in each of the clusters.** Number of cells for each of the major cell types in our dataset. It further gives the number of cells present in each cluster of the cell type.

**Table S8.**

**A) CD4 T cells**

| gene | p_val | avg_logFC | pct.1 | pct.2 | p_val_adj | cluster |
|---|---|---|---|---|---|---|
| **FGFBP2** | 5.08E-90 | 1.3788028 | 0.164 | 0.043 | 2.46E-87 | 1 |
| **NKG7** | 3.79E-109 | 1.3413716 | 0.294 | 0.112 | 1.84E-106 | 1 |
| **GZMH** | 2.01E-85 | 1.2617363 | 0.16 | 0.043 | 9.77E-83 | 1 |
| **GZMA** | 6.90E-62 | 1.0378083 | 0.191 | 0.073 | 3.35E-59 | 1 |
| **GNLY** | 9.54E-97 | 1.0227902 | 0.264 | 0.097 | 4.63E-94 | 1 |
| **CXCR5** | 3.82E-127 | 1.707846 | 0.168 | 0.013 | 1.85E-124 | 7 |
| **CHI3L2** | 1.84E-28 | 0.9722734 | 0.102 | 0.027 | 8.93E-26 | 8 |
| **GZMH** | 8.42E-84 | 1.9580125 | 0.349 | 0.058 | 4.08E-81 | 9 |
| **GNLY** | 4.08E-69 | 1.8397658 | 0.465 | 0.12 | 1.98E-66 | 9 |
| **S100A9** | 9.49E-18 | 1.8062813 | 0.201 | 0.068 | 4.60E-15 | 9 |
| **NKG7** | 4.79E-67 | 1.6259264 | 0.498 | 0.137 | 2.33E-64 | 9 |
| **CCL3** | 9.86E-17 | 1.5112569 | 0.152 | 0.044 | 4.78E-14 | 9 |
| **FGFBP2** | 8.98E-62 | 1.308841 | 0.316 | 0.06 | 4.36E-59 | 9 |
| **FCN1** | 1.20E-15 | 1.2569039 | 0.134 | 0.038 | 5.83E-13 | 9 |
| **GZMA** | 7.07E-24 | 0.8354358 | 0.279 | 0.091 | 3.43E-21 | 9 |
| **TBX21** | 1.85E-22 | 0.8232073 | 0.141 | 0.031 | 9.00E-20 | 9 |
| **HOPX** | 5.70E-23 | 0.7992697 | 0.245 | 0.077 | 2.76E-20 | 9 |
| **GZMB** | 9.29E-16 | 0.6872567 | 0.134 | 0.037 | 4.51E-13 | 9 |
| **IFNG** | 1.67E-14 | 0.6800668 | 0.145 | 0.044 | 8.12E-12 | 9 |
| **PRF1** | 8.95E-12 | 0.6017679 | 0.104 | 0.03 | 4.34E-09 | 9 |
| **GZMK** | 1.88E-50 | 1.5009135 | 0.22 | 0.039 | 9.11E-48 | 10 |
| **CCL5** | 6.47E-54 | 1.2063735 | 0.622 | 0.241 | 3.14E-51 | 10 |
| **CST7** | 5.40E-39 | 1.1132755 | 0.458 | 0.166 | 2.62E-36 | 10 |
| **CXCR3** | 5.55E-27 | 1.0689779 | 0.168 | 0.039 | 2.69E-24 | 10 |
| **GZMA** | 6.93E-25 | 0.9999948 | 0.273 | 0.091 | 3.36E-22 | 10 |
| **HOPX** | 1.14E-17 | 0.8881745 | 0.217 | 0.077 | 5.54E-15 | 10 |
| **NKG7** | 5.83E-27 | 0.7453275 | 0.374 | 0.14 | 2.83E-24 | 10 |
| **TBX21** | 2.77E-12 | 0.6945313 | 0.108 | 0.032 | 1.35E-09 | 10 |

| | | | | | | |
|---|---|---|---|---|---|---|
| **IFNG** | 3.66E-13 | 0.4985953 | 0.136 | 0.044 | 1.78E-10 | 10 |
| **GNLY** | 0 | 2.6907898 | 0.802 | 0.108 | 0 | 12 |
| **GZMB** | 6.27E-235 | 2.2753367 | 0.383 | 0.029 | 3.04E-232 | 12 |
| **NKG7** | 1.08E-276 | 2.2584279 | 0.793 | 0.126 | 5.24E-274 | 12 |
| **FGFBP2** | 6.18E-268 | 2.1633731 | 0.538 | 0.051 | 3.00E-265 | 12 |
| **CCL5** | 1.68E-185 | 1.8882198 | 0.857 | 0.232 | 8.17E-183 | 12 |
| **GZMH** | 5.01E-187 | 1.8078891 | 0.456 | 0.053 | 2.43E-184 | 12 |
| **FCGR3A** | 5.81E-28 | 1.6992762 | 0.103 | 0.018 | 2.82E-25 | 12 |
| **CCL4** | 1.17E-48 | 1.5732819 | 0.167 | 0.026 | 5.69E-46 | 12 |
| **PRF1** | 7.00E-66 | 1.5444357 | 0.195 | 0.027 | 3.39E-63 | 12 |
| **CTSW** | 2.44E-58 | 1.5125388 | 0.356 | 0.09 | 1.18E-55 | 12 |
| **CST7** | 6.83E-129 | 1.4397931 | 0.663 | 0.158 | 3.31E-126 | 12 |
| **HOPX** | 1.24E-85 | 1.3983094 | 0.374 | 0.072 | 6.04E-83 | 12 |
| **ITGAM** | 1.10E-66 | 1.3803749 | 0.167 | 0.02 | 5.35E-64 | 12 |
| **KLRC4.KLRK1** | 7.69E-80 | 1.3773626 | 0.17 | 0.017 | 3.73E-77 | 12 |
| **KLRK1** | 7.60E-75 | 1.3567504 | 0.164 | 0.017 | 3.68E-72 | 12 |
| **KLRF** | 1.68E-57 | 1.1605099 | 0.103 | 0.008 | 8.16E-55 | 12 |
| **GZMA** | 6.48E-54 | 1.129899 | 0.343 | 0.088 | 3.14E-51 | 12 |
| **TARP-refseq** | 1.24E-47 | 1.1269784 | 0.252 | 0.055 | 6.01E-45 | 12 |
| **CCL3** | 9.26E-25 | 1.0930488 | 0.164 | 0.043 | 4.49E-22 | 12 |
| **FOXP3** | 9.99E-58 | 1.8722019 | 0.171 | 0.017 | 4.85E-55 | 13 |
| **LGALS3** | 3.63E-26 | 1.3802594 | 0.251 | 0.065 | 1.76E-23 | 13 |
| **CTLA4** | 1.45E-14 | 0.8787505 | 0.209 | 0.069 | 7.05E-12 | 13 |
| **CXCR5** | 2.26E-99 | 1.7331973 | 0.176 | 0.014 | 1.10E-96 | 14 |
| **CXCR3** | 1.15E-08 | 0.6471964 | 0.104 | 0.04 | 5.59E-06 | 14 |
| **GZMK** | 8.74E-19 | 1.0145717 | 0.123 | 0.04 | 4.24E-16 | 15 |
| **CXCR3** | 1.16E-11 | 0.7324178 | 0.103 | 0.039 | 5.60E-09 | 15 |
| **CXCR3** | 2.54E-14 | 0.8116743 | 0.112 | 0.039 | 1.23E-11 | 16 |

**B) CD8 T cells**

| gene | p_val | avg_logFC | pct.1 | pct.2 | p_val_adj | cluster |
|---|---|---|---|---|---|---|
| **LEF1** | 0 | 1.8988998 | 0.523 | 0.091 | 0 | 2 |
| **CCR7** | 2.50E-307 | 1.7167545 | 0.347 | 0.057 | 1.21E-304 | 2 |
| **PASK** | 7.85E-214 | 1.5544292 | 0.34 | 0.08 | 3.81E-211 | 2 |
| **MYC** | 1.02E-190 | 1.5165717 | 0.347 | 0.092 | 4.93E-188 | 2 |
| **SELL** | 5.72E-207 | 1.4364113 | 0.402 | 0.116 | 2.78E-204 | 2 |
| **TXK** | 2.03E-127 | 1.3364083 | 0.142 | 0.021 | 9.85E-125 | 2 |
| **CD27** | 3.15E-119 | 1.2886679 | 0.273 | 0.084 | 1.53E-116 | 2 |
| **IL7R** | 2.52E-155 | 1.1706575 | 0.429 | 0.161 | 1.22E-152 | 2 |
| **FOXP1** | 1.27E-36 | 0.8811065 | 0.12 | 0.045 | 6.16E-34 | 2 |
| **MYC** | 7.33E-75 | 1.3937987 | 0.345 | 0.111 | 3.56E-72 | 6 |
| **IL4R** | 7.05E-50 | 1.0419241 | 0.239 | 0.076 | 3.42E-47 | 6 |
| **CD28** | 1.07E-37 | 0.8430301 | 0.114 | 0.026 | 5.19E-35 | 6 |
| **BIRC3** | 1.60E-26 | 0.8148054 | 0.133 | 0.043 | 7.76E-24 | 6 |
| **ICOS** | 2.66E-20 | 0.704204 | 0.123 | 0.044 | 1.29E-17 | 6 |
| **CCR7** | 2.00E-29 | 0.6153928 | 0.218 | 0.085 | 9.71E-27 | 6 |
| **KLRF1** | 3.88E-40 | 0.9667987 | 0.126 | 0.036 | 1.88E-37 | 7 |
| **S100A9** | 4.10E-11 | 1.5853778 | 0.187 | 0.07 | 1.99E-08 | 8 |
| **FCN1** | 8.60E-07 | 0.6327533 | 0.101 | 0.035 | 0.000417 | 8 |
| **LEF1** | 1.17E-119 | 1.3393149 | 0.426 | 0.126 | 5.66E-117 | 9 |
| **CCR7** | 1.06E-75 | 1.3370251 | 0.281 | 0.081 | 5.16E-73 | 9 |
| **PASK** | 1.53E-60 | 1.1976862 | 0.293 | 0.101 | 7.42E-58 | 9 |
| **SELL** | 3.10E-71 | 1.0802719 | 0.372 | 0.137 | 1.51E-68 | 9 |
| **MYC** | 6.62E-64 | 0.9356279 | 0.319 | 0.111 | 3.21E-61 | 9 |
| **TCF7** | 3.21E-23 | 0.9024087 | 0.142 | 0.055 | 1.56E-20 | 9 |
| **TRDC** | 6.11E-61 | 1.741988 | 0.138 | 0.02 | 2.96E-58 | 12 |
| **FCGR3A** | 6.71E-97 | 1.4408549 | 0.394 | 0.096 | 3.26E-94 | 12 |
| **KLRC3** | 1.69E-43 | 1.2377133 | 0.181 | 0.044 | 8.18E-41 | 12 |
| **GZMB** | 1.95E-101 | 1.2367893 | 0.594 | 0.193 | 9.45E-99 | 12 |
| **KLRC1** | 3.44E-59 | 1.2042799 | 0.167 | 0.029 | 1.67E-56 | 12 |

| | | | | | | |
|---|---|---|---|---|---|---|
| **KLRB1** | 1.43E-50 | 1.0618519 | 0.379 | 0.139 | 6.96E-48 | 12 |
| **LYN** | 1.29E-27 | 0.9356006 | 0.138 | 0.037 | 6.27E-25 | 12 |
| **KLRF1** | 4.69E-37 | 0.8531752 | 0.158 | 0.038 | 2.27E-34 | 12 |
| **S1PR5** | 8.45E-31 | 0.7701552 | 0.271 | 0.103 | 4.10E-28 | 12 |
| **TTC38** | 3.23E-16 | 0.6734551 | 0.119 | 0.041 | 1.56E-13 | 12 |
| **ITGAX** | 1.36E-128 | 2.1035366 | 0.394 | 0.029 | 6.57E-126 | 13 |
| **TRDC** | 5.23E-94 | 1.8934453 | 0.289 | 0.021 | 2.54E-91 | 13 |
| **KLRC3** | 3.44E-40 | 1.3505882 | 0.289 | 0.046 | 1.67E-37 | 13 |
| **LYN** | 1.04E-14 | 0.993765 | 0.169 | 0.039 | 5.03E-12 | 13 |
| **CD160** | 1.43E-23 | 0.9535271 | 0.232 | 0.048 | 6.91E-21 | 13 |
| **FCGR3A** | 1.86E-16 | 0.9088716 | 0.324 | 0.105 | 9.01E-14 | 13 |
| **PIK3AP1** | 2.55E-23 | 0.8608042 | 0.31 | 0.077 | 1.24E-20 | 13 |
| **KLRC1** | 1.29E-13 | 0.7619082 | 0.148 | 0.033 | 6.27E-11 | 13 |

**C) Classical monocytes**

| gene | p_val | avg_logFC | pct.1 | pct.2 | p_val_adj | cluster |
|---|---|---|---|---|---|---|
| **FCER1A** | 3.49E-158 | 2.397606 | 0.205 | 0.006 | 1.69E-155 | 5 |
| **CLEC10A** | 2.67E-57 | 1.945571 | 0.287 | 0.076 | 1.29E-54 | 5 |
| **CD1C** | 1.08E-28 | 1.199427 | 0.145 | 0.036 | 5.23E-26 | 5 |

**D) Intermediate monocytes**

| gene | p_val | avg_logFC | pct.1 | pct.2 | p_val_adj | cluster |
|---|---|---|---|---|---|---|
| **S100A12** | 2.36E-26 | 1.4033592 | 0.455 | 0.137 | 1.15E-23 | 3 |
| **CD14** | 7.58E-27 | 1.0214426 | 0.57 | 0.22 | 3.68E-24 | 3 |
| **CD163** | 6.63E-17 | 0.718982 | 0.223 | 0.046 | 3.22E-14 | 3 |
| **CLEC4E** | 1.15E-14 | 0.7058619 | 0.293 | 0.091 | 5.59E-12 | 3 |
| **THBS1** | 3.05E-13 | 0.6305201 | 0.347 | 0.134 | 1.48E-10 | 3 |
| **MGST1** | 1.16E-12 | 0.5038258 | 0.182 | 0.042 | 5.62E-10 | 3 |
| **RNASE2** | 2.01E-11 | 0.4517372 | 0.174 | 0.043 | 9.73E-09 | 3 |
| **RNASE6** | 1.36E-10 | 0.451708 | 0.26 | 0.095 | 6.57E-08 | 3 |

**E) B cells**

| gene | p_val | avg_logFC | pct.1 | pct.2 | p_val_adj | cluster |
|---|---|---|---|---|---|---|
| **IGHA1-secreted** | 2.01E-61 | 1.669628789 | 0.397 | 0.094 | 9.77E-59 | 2 |
| **CD27** | 2.85E-21 | 0.880595753 | 0.135 | 0.029 | 1.38E-18 | 2 |
| **ITGAX** | 9.56E-32 | 1.286536823 | 0.236 | 0.038 | 4.64E-29 | 5 |
| **CST7** | 2.41E-14 | 1.229898092 | 0.112 | 0.02 | 1.17E-11 | 5 |
| **IGHG2-secreted** | 9.55E-52 | 1.133017096 | 0.742 | 0.249 | 4.63E-49 | 5 |
| **IGHG1-secreted** | 1.18E-46 | 1.058429191 | 0.646 | 0.199 | 5.71E-44 | 5 |
| **NKG7** | 6.77E-11 | 0.988539041 | 0.163 | 0.049 | 3.28E-08 | 5 |
| **LILRB1** | 3.44E-06 | 0.819106799 | 0.107 | 0.036 | 0.001669 | 5 |
| **IGHG3-secreted** | 8.80E-09 | 0.713353526 | 0.129 | 0.038 | 4.27E-06 | 5 |
| **FCGR2A** | 1.61E-12 | 0.681229262 | 0.112 | 0.022 | 7.80E-10 | 5 |
| **IGHG1-secreted** | 1.63E-75 | 1.431057298 | 0.635 | 0.175 | 7.92E-73 | 6 |
| **IGHG2-secreted** | 7.83E-65 | 1.347913711 | 0.681 | 0.228 | 3.80E-62 | 6 |
| **CD27** | 1.10E-20 | 0.868503278 | 0.141 | 0.03 | 5.31E-18 | 6 |
| **IGHG3-secreted** | 2.96E-09 | 0.865141203 | 0.107 | 0.036 | 1.44E-06 | 6 |
| **IGHG4-secreted** | 5.03E-24 | 0.659245382 | 0.104 | 0.013 | 2.44E-21 | 6 |
| **NCR3** | 6.32E-13 | 0.519980657 | 0.15 | 0.049 | 3.07E-10 | 6 |
| **CD86** | 2.18E-13 | 0.516502142 | 0.138 | 0.041 | 1.06E-10 | 6 |
| **CTSA** | 2.91E-10 | 0.434407395 | 0.153 | 0.058 | 1.41E-07 | 6 |

**Table S8. Significantly differentially expressed genes for each cell type (A-E).** Differentially expressed genes are from one cluster compared against the rest in the same cell type. gene: gene name, p_val: raw p value, avg_logFC: average log2 fold change, pct.1: percent of cells in hat cluster that express this gene, pct.2: percent of cells in all other clusters that express this gene, p_val_adj: p value adjusted by Benjamini-Hochberg for multiple comparisons, cluster: cluster number as noted in UMAP. Significant genes were defined as adjusted $p<0.05$, avg_locFC>0, and pct1/pct2>2.5.

**Table S9.**

| | CD4 T cells | CD8 T cells | Classical monocytes | Intermediate monocytes | Nonclassical monocytes | B cells | NK cells |
|---|---|---|---|---|---|---|---|
| 1 | IL32 | IL32 | CCL3 | IL32 | CD52 | CD74 | IL32 |
| 2 | KLF2 | BTG1 | CCL4 | KLF2 | FCN1 | CD83 | KLRB1 |
| 3 | SELL | CD52 | IL1B | SELL | LGALS1 | CD79B | GNLY |
| 4 | RGS1 | KLF2 | TNF | RGS1 | CD74 | IGHM-membrane | GZMA |
| 5 | CCR7 | TRAC | DUSP2 | CCR7 | TSPAN14 | IGHD-membrane | CCL5 |
| 6 | SELPLG | HLA.B | IFITM3 | SELPLG | CCL4 | TCL1A | SELPLG |
| 7 | IFITM3 | HOPX | MNDA | IFITM3 | PTPRC | R3HDM4 | FCER1G |
| 8 | GIMAP5 | GNLY | IER3 | GIMAP5 | IL6 | KLF2 | BTG1 |
| 9 | CD69 | CCL3 | IL6 | CD69 | ITGB2 | CD79A | CD52 |
| 10 | TNFSF10 | CXCR4 | SOD2 | TCF7 | C10orf54 | SLC2A3 | CXCR4 |

**Table S9. Top 10 disease-regulated genes for each cell type.**

**Table S10.**

| Patient number | Pat_Type | Total no. of cells | B cells | CD4 T cells | CD8 T cells | Monocytes | NK cells |
|---|---|---|---|---|---|---|---|
| 1 | CRT | 1658 | 29 | 310 | 1150 | 129 | 40 |
| 2 | C | 1869 | 326 | 791 | 465 | 263 | 24 |
| 3 | H | 1799 | 119 | 277 | 1157 | 151 | 95 |
| 4 | H- | 1164 | 112 | 616 | 307 | 115 | 14 |
| 5 | CRT | 358 | 22 | 91 | 205 | 11 | 29 |
| 7 | H | 2574 | 106 | 603 | 951 | 683 | 231 |
| 8 | H- | 394 | 40 | 152 | 65 | 91 | 46 |
| 9 | CRT | 708 | 98 | 210 | 241 | 98 | 61 |
| 10 | C | 951 | 87 | 291 | 432 | 110 | 31 |
| 11 | H | 514 | 32 | 141 | 144 | 179 | 18 |
| 12 | H- | 675 | 22 | 292 | 194 | 159 | 8 |
| 13 | CRT | 964 | 68 | 187 | 415 | 198 | 96 |
| 14 | C | 737 | 51 | 71 | 486 | 113 | 16 |
| 15 | H | 1366 | 43 | 207 | 294 | 788 | 34 |
| 16 | H- | 1266 | 60 | 697 | 265 | 160 | 84 |
| 17 | CRT | 748 | 90 | 202 | 259 | 161 | 36 |
| 18 | C | 1178 | 70 | 414 | 516 | 144 | 34 |
| 19 | H | 1323 | 98 | 527 | 465 | 195 | 38 |
| 20 | H- | 1244 | 66 | 801 | 174 | 150 | 53 |
| 21 | CRT | 973 | 145 | 254 | 289 | 226 | 59 |
| 22 | C | 1836 | 157 | 179 | 985 | 455 | 60 |
| 23 | H | 958 | 50 | 77 | 486 | 203 | 142 |
| 24 | H- | 1094 | 69 | 488 | 347 | 168 | 22 |
| 25 | CRT | 1436 | 164 | 559 | 274 | 367 | 72 |
| 26 | C | 1141 | 72 | 290 | 506 | 97 | 176 |
| 27 | H | 766 | 112 | 350 | 186 | 101 | 17 |
| 28 | H- | 964 | 70 | 427 | 222 | 195 | 50 |
| 29 | CRT | 1017 | 56 | 218 | 421 | 282 | 40 |

| | | | | | | | |
|---|---|---|---|---|---|---|---|
| 30 | C | 1117 | 217 | 300 | 213 | 248 | 139 |
| 31 | H | 961 | 101 | 350 | 262 | 199 | 49 |
| 32 | H- | 1526 | 167 | 673 | 467 | 190 | 29 |

**Table S10: Number of Cells for each participant.** Number of cells for each Patient in our dataset. Patient 6 has no cells. The column Pat_Type gives the type of the patient. 'CRT' denotes a patient that is HIV+, CVD+, cholesterol-reducing therapy+. C denotes a patient that is HIV+, CVD+, cholesterol-reducing therapy-. H denotes a patient that is HIV+, CVD-, cholesterol-reducing therapy-. H- denotes a patient that is HIV-, CVD-, cholesterol-reducing therapy

**Table S11.**

| Antibody | Gene | All | CD4T | CD8T | CM | INT | NCM | B | NK |
|---|---|---|---|---|---|---|---|---|---|
| CD11b (ITGAM) | ITGAM | 0 | 0 | 0 | 0 | 0 | 0 | 0 | 0.1466 |
| CD11c (ITGAX) | ITGAX | 0 | 0 | 0 | 0 | 0.0109 | 0 | 0 | 0.066 |
| CD123 (IL3RA) | IL3RA | 0.1749 | 0 | 0 | 0.0469 | 0.3161 | 0 | 0 | 0 |
| CD127 (IL7R) | IL7R | 0.1008 | 0.0809 | 0.0906 | 0 | 0 | 0 | 0 | 0 |
| CD137 (TNFRSF9) | TNFRSF9 | 0 | 0 | 0 | 0 | 0 | 0 | 0 | 0 |
| CD141 (THBD) | THBD | 0 | 0 | 0 | 0 | 0 | 0 | 0 | 0 |
| CD14 | CD14 | 0 | 0 | 0 | 0 | 0.275 | 0 | 0 | 0 |
| CD152 (CTLA4) | CTLA4 | 0 | 0 | 0 | 0 | 0 | 0 | 0 | 0 |
| CD163 | CD163. PolyA1 | 0 | 0 | 0 | 0 | 0 | 0 | 0 | 0 |
| CD16 (FCGR3A) | FCGR3A | 0.3022 | 0 | 0.0987 | 0 | 0.4154 | 0.0932 | 0 | 0 |
| CD183 (CXCR3) | CXCR3 | 0.0021 | 0.031 | 0.0533 | 0 | 0 | 0 | 0 | 0 |
| CD184 (CXCR4) | CXCR4 | 0 | 0.0615 | 0.1092 | 0 | 0.0123 | 0.0822 | 0.1891 | 0.1222 |
| CD185 (CXCR5) | CXCR5 | 0 | 0 | 0 | 0 | 0 | 0 | 0 | 0 |
| CD192 | CCR2 | 0 | 0 | 0 | 0 | 0 | 0 | 0 | 0 |
| CD194 (CCR4) | CCR4 | 0 | 0 | 0 | 0 | 0 | 0 | 0 | 0 |

| | | | | | | | | | |
|---|---|---|---|---|---|---|---|---|---|
| CD195 (CCR5) | CCR5 | 0 | 0 | 0 | 0 | 0 | 0 | 0 | 0 |
| CD196 (CCR6) | CCR6. PolyA1 | 0 | 0 | 0 | 0 | 0 | 0 | 0 | 0 |
| CD196 (CCR6) | CCR6 | 0 | 0 | 0 | 0 | 0 | 0 | 0 | 0 |
| CD197 (CCR7) | CCR7 | 0 | 0 | 0 | 0 | 0 | 0 | 0 | 0 |
| CD20 (MS4A1) | MS4A1. PolyA1 | 0.0968 | 0 | 0 | 0 | 0 | 0 | 0.0865 | 0 |
| CD223 (LAG3) | LAG3 | 0 | 0 | 0 | 0 | 0 | 0 | 0 | 0 |
| CD25 (IL2RA) | IL2RA | 0.0541 | 0.0813 | 0 | 0 | 0 | 0 | 0 | 0 |
| CD27 | CD27 | 0.0947 | 0 | 0.2028 | 0 | 0 | 0 | 0 | 0 |
| CD2 | CD2. PolyA1 | 0 | 0 | 0 | 0 | 0 | 0 | 0 | 0 |
| CD2 | CD2 | 0 | 0 | 0 | 0 | 0 | 0 | 0 | 0.0117 |
| CD36 | CD36 | 0 | 0 | 0 | 0 | 0.1147 | 0 | 0 | 0 |
| CD38 | CD38 | 0 | 0 | 0 | 0 | 0 | 0 | 0 | 0 |
| CD3 (CD3E) | CD3D | 0.0056 | 0.041 | 0 | 0 | 0 | 0 | 0 | 0 |
| CD3 (CD3E) | CD3E | 0 | 0.0421 | 0 | 0 | 0 | 0 | 0 | 0 |
| CD3 (CD3E) | CD3G. PolyA1 | 0 | 0 | 0 | 0 | 0 | 0 | 0 | 0 |
| CD45RA (PTPRC) | PTPRC. PolyA1 | 0 | 0 | 0 | 0 | 0.1042 | 0 | 0 | 0 |
| CD45RO (PTPRC) | PTPRC. PolyA1 | 0.1011 | 0 | 0 | 0 | 0 | 0 | 0 | 0 |

| | | | | | | | | | |
|---|---|---|---|---|---|---|---|---|---|
| CD4 | CD4 | 0.2776 | 0 | 0 | 0.0589 | 0.0245 | 0 | 0 | 0 |
| CD56 (NCAM1) | NCAM1 | 0 | 0 | 0 | 0 | 0 | 0 | 0 | 0 |
| CD69 | CD69 | 0 | 0.0311 | 0 | 0 | 0 | 0 | 0 | 0 |
| CD86 | CD86. PolyA1 | 0 | 0 | 0 | 0 | 0 | 0 | 0 | 0 |
| CD8 (CD8A) | CD8A | 0.0408 | 0 | 0.0264 | 0 | 0 | 0 | 0 | 0 |
| CD8 (CD8A) | CD8B | 0 | 0 | 0 | 0 | 0 | 0 | 0 | 0 |
| CD9 | CD9 | 0 | 0 | 0 | 0 | 0 | 0 | 0 | 0 |
| HLA.DR (CD74) | CD74 | 0.4199 | 0.2219 | 0.2303 | 0.3737 | 0.5002 | 0.2321 | 0.0864 | 0.215 |
| HLA.DR (CD74) | HLA.DRA | 0.4471 | 0.0987 | 0.2437 | 0.4888 | 0.5097 | 0.3102 | 0.1178 | 0 |

**Table S11. Non-negative Spearman correlation between antibody and gene in each cell type.** All; all cell types, CM; Classical monocytes, INT; intermediate monocytes, NCM; Nonclassical monocytes. Dense color indicates higher value.

**Data S1. (separate excel file)**

Data underlying Figure 4 dotplots. Average gene expression per cell in each of the clusters for the main subsets: CD4, CD8 T cells, Classical, Intermediate, Non-classical monocytes, B cells and NK cells.

**Data S2. (separate excel file)**

Data underlying Figure 6 volcano plots. Differentially expressed genes [HIV-CVD- vs HIV+CVD-, HIV-CVD+ vs HIV+CVD+, HIV+CVD+ vs HIV+CVD+CRT (cholesterol reducing treatment)] compared in each cell cluster.
